# A Mediterranean-like fat blend protects against the development of severe colitis in the mucin-2 deficient murine model

**DOI:** 10.1101/2021.09.27.462042

**Authors:** Natasha Haskey, Jiayu Ye, Mehrbod Estaki, Andrea A. Verdugo Meza, Jacqueline A. Barnett, Mitra Yousefi, Blake W. Birnie, Samantha Gruenheid, Sanjoy Ghosh, Deanna L. Gibson

## Abstract

The Mediterranean diet (MD) is a health-promoting, moderate fat diet (~ 40% energy). It is not known if the blend of fats found in the MD contribute to the beneficial protective effects. We compared the MD fat blend (high monounsaturated, 2:1 n-6:n-3 polyunsaturated and moderate saturated fat) to isocaloric diets composed with corn oil (CO, n-6 polyunsaturated-rich), olive oil (high monounsaturated-rich) or milk fat (MF, saturated-rich) on spontaneous colitis development in Muc2^−/−^ mice. The MD resulted in lower clinical and histopathological scores, and induced tolerogenic CD103+CD11b+ dendritic, Th22 and IL-17+IL-22+ cells important for intestinal barrier repair. MD also reduced attendant insulin resistance and a shift to a higher health-promoting gut microbes including *Lactobacillus B. animalis* and Muribaculaceae, whereas CO showed higher prevalence of mucin-degraders (*Akkermansia muciniphila)* and colitis promoters (Enterobacteriaceae). Our findings suggest that the MD fat blend could be recommended as a maintenance diet for colitis.

## 1. Introduction

Inflammatory bowel diseases (IBD), including both ulcerative colitis (UC) and Crohn’s disease (CD) are characterized by chronic intestinal inflammation, along with detrimental changes to the gut microbiome which can lead to broader systemic effects, including metabolic abnormalities (reviewed in Verdugo-Meza et al., 2020). The complex relationship between diet, genetics, environment, and alterations in the gut microbiome are known etiological factors involved in the development of IBD (Ramos and Papadakis, 2019). The rapid increase in the incidence and prevalence of this chronic condition in continents such as Asia, Africa, South America, and Eastern Europe could point to the important role diet plays in disease development (Ng *et al.*, 2017). In particular, the increased consumption of a Western diet pattern (WD) characterized by high n-6 polyunsaturated (n-6 PUFA) vegetable oils, processed meat, sweetened beverages, environmental contaminants, and food additives with a concomitant reduction in protective phytochemicals fibre, fruits, vegetables, and fish is associated with an increased risk of developing UC along with other chronic inflammatory diseases (Wong *et al.*, 2015; Barnett and Gibson, 2020; T. Li *et al.*, 2020). Thereby, it appears that, the WD promotes local and systemic inflammation driven by changes in gut microbiota and the immune system, affecting the gut integrity (Christ, Lauterbach and Latz, 2019).

Much of the research on diet and IBD has focused on the negative impact of high dietary fat intake and its association with IBD (Tjonneland *et al.*, 2009; Hou, Abraham and El-Serag, 2011; Ananthakrishnan *et al.*, 2014). However, one high fat diet (40% by energy) that has been associated with beneficial effects in immune and metabolic diseases including IBD is the Mediterranean diet (MD) (Sasson et al., 2021). Additionally, we and others have shown that the type of fat, independent of caloric content, influence intestinal inflammation, metabolism, and host-microbe function (Patterson et al., 2014; DeCoffe et al., 2016; Abulizi et al., 2019). For example, murine models have demonstrated that n-6 PUFA and saturated fatty acids (SFA) result in inflammation-induced colonic damage while monounsaturated fatty acids (MUFA) are protective. Additionally, the benefits of n-3 PUFA, commonly present in fish oils may be dependent on SFA in the diet (DeCoffe *et al.*, 2016). SFA derived from dairy fat are unique in their compensatory inflammatory responses involved in tissue restitution (DeCoffe et al., 2016; Abulizi et al., 2019). Human observational studies show that after energy-adjustment for total fat intake, SFA, such as myristic acid and long-term intake of trans- fatty acids and n-6 PUFA, are associated with an increasing incidence and risk of flare-up in UC patients (Tjonneland *et al.*, 2009; Ananthakrishnan *et al.*, 2014). Understanding the mechanisms of how various fatty acids impact IBD is important in the development of evidence-based guidelines to reduce specific food-induced inflammation, promote remission and dietary tolerance.

While nutrition consensus guidelines for IBD are lacking, a review of clinical studies recommend a diet rich in fruits and vegetables, n-3 PUFA and low n-6 PUFA as a maintenance diet for IBD (Forbes *et al.*, 2017; Haskey and Gibson, 2017). A specific dietary pattern for the management of IBD is yet to be defined. Recent evidence suggests that the MD could be a potential diet strategy due to its beneficial anti-inflammatory and antioxidant properties that have been documented against other chronic diseases with inflammatory origin (Bloomfield *et al.*, 2016; Sasson *et al.*, 2021) and the mechanisms underlying the potential benefits of this diet both active and chronic colitis are lacking. More specifically, the precise impact of the blend of fats contained in the MD on intestinal inflammation and the microbiome have not been elucidated.

Therefore, the primary aim of this study was to compare a Mediterranean-like fat blend (40.8% calories from fat with 7.8% SFA, 27.7% MUFA and 4.5% total PUFA (0.8% derived from n-3 PUFA)) to fat from corn oil (CO), milk fat (MF), and olive oil (OO) on intestinal inflammation, aspects of lipid and glucose metabolism, and the gut microbiome in a spontaneous murine model of colitis. An experimental model of spontaneous and chronic colitis, the mucin 2 knock-out mouse strain (Muc2^−/−^) was used to mimic human UC. Mice that lack mucin 2 demonstrate loss and function of goblet cells that produce the main secretory gel-forming mucin (Muc2) found in the lumen of the colon which result in rectal prolapse and damage driven by RELM-β (Morampudi *et al.*, 2016), as well as develop metabolic co-morbidities (Ye *et al.*, 2021). Overall, we show that Muc2^−/−^ mice fed the MD were protected from the development of severe colitis and impairments in glucose tolerance through anti-inflammatory host defence mechanisms. Uniquely, the MD had a higher ratio of ASVs of health-promoting microbes such as *Lactobacillus B. animalis* and Muribaculaceae.

## 2. Results

### MD protects against severe clinical disease activity, histological damage and promotes intestinal homeostasis

To understand the effect of the MD fat on colitis, we fed Muc2^−/−^ mice a 40.8% fat diet blend (27.7% MUFA, 7.8% SFA, and 4.5% total PUFA (n-6 PUFA to n-3 PUFA; 2:1) for nine weeks (12 weeks of age), compared to isocaloric and isonitrogenous diets altered in fat only (corn oil rich in n-6 PUFA, olive oil rich in MUFA and milk fat rich in SFA) (**Table 1**). The total fat content is nutritionally relevant because it mimics the total fat content of the Mediterranean diet consumed by humans (Bach-Faig *et al.*, 2011). No differences in food intake or weight changes were seen between any of the diet groups **(Figure S1A and S1B)**, which indicates that variations in colitis was not driven by increased caloric intake or weight gain. Mice fed the MD and OO diets had significantly lower disease activity index (DAI) scores than the MF diet, with the MD showing the lowest DAI in comparison to the other diet groups (**Figure 1A and S1C)**. Ten percent of the CO and MF mice developed rectal prolapse, the most severe form of colitis assessed by our scoring system. No rectal prolapses were seen in the MD or OO diets (**Figure 1C**). The CO diet had an earlier onset of rectal prolapse (week 6, week 7 and week 9) than the MF diet (week 8 and 2 mice at week 9) suggesting the mice fed CO developed a more aggressive form of colitis earlier. Shortening of the colon, a hallmark sign of increased inflammation and disease severity in colitis (Kim *et al.*, 2012), was not seen in the MD or MF fed diet, whereas the CO and OO fed diets had significantly shorter colons than mice fed the MD (**Figure 1B**). In agreement with the clinical observations, histopathological differences were observed between the diet groups (**Figure 1D to 1I)** with crypt hyperplasia, edema and a loss of epithelial integrity and inflammatory changes seen in all groups consistent with the model (Morampudi *et al.*, 2016). The MD group showed the lowest total histopathological scores with the least epithelial damage, in addition to less ulcerations and abscesses (**Figure 1D to 1I)**. Overall, the Muc2^−/−^ mice fed the CO and MF diet exhibited multiple features of a more aggressive colitis, whereas the MD and OO diet presented with a milder form of disease.

**Figure 1.**
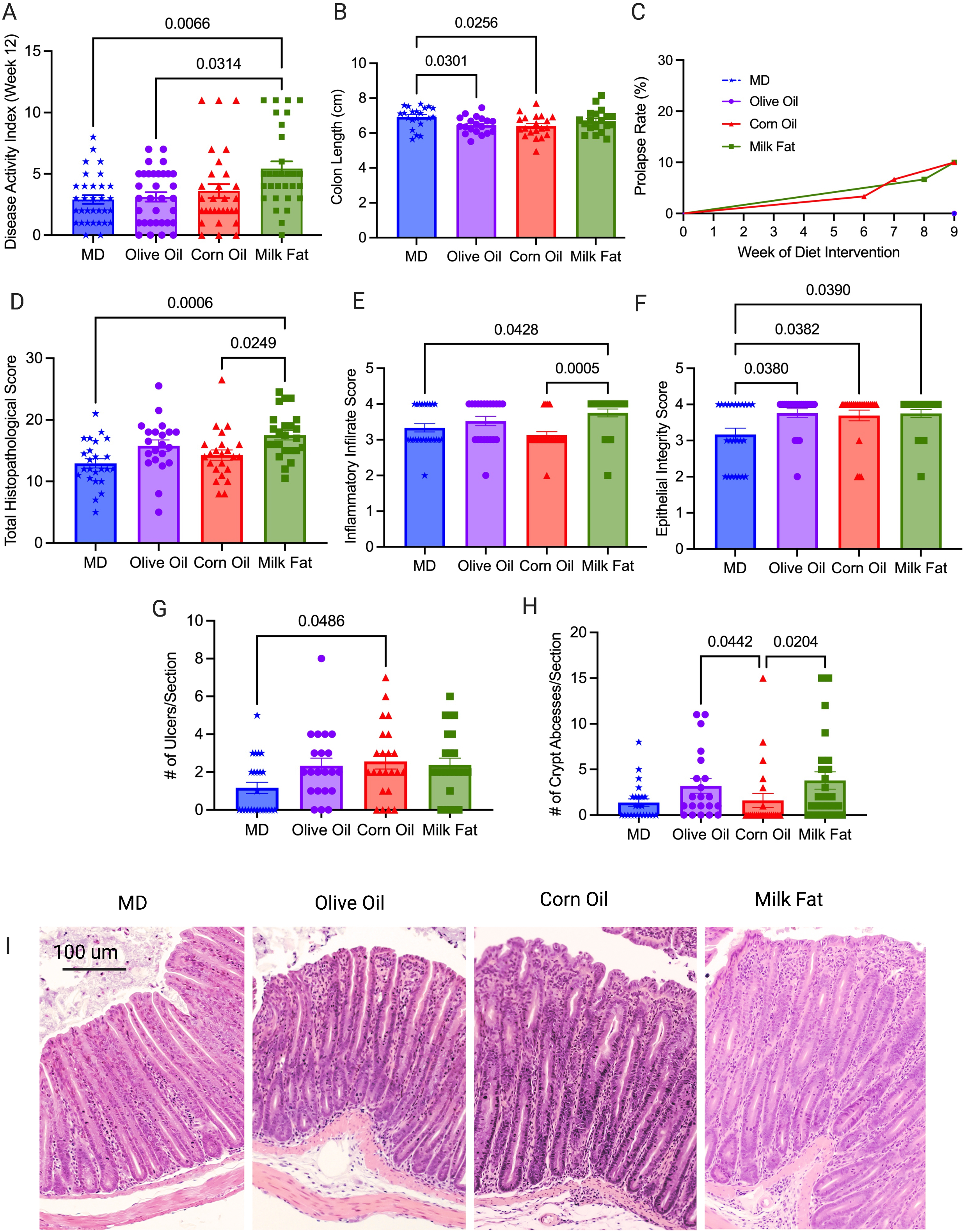
MD protects against severe clinical disease activity and histological damage. Colitis severity was measured by (A) disease activity index at endpoint in four diet groups: MD, olive oil, corn oil and milk fat diets, (B) colon length at endpoint, (C) rectal prolapse rate of the mice, (D) total histopathological scores, (E) inflammatory infiltrate score (F) epithelial integrity scores (G) ulcers per tissue section and (H) crypt abscesses per tissue section (I) representative H & E stained cross sections of the distal colon (images taken at 10x magnification). Colons analysed in (D) were scored for inflammation, ulceration, edema, epithelial integrity, and hyperplasia. *n*=30-33 mice/group. Statistical significance was determined by Kruskal-Wallis with Dunn’s post-hoc analysis, means ± SEM. MD: Mediterranean diet; SEM: standard error of mean. *p*<0.05. (See also Figure S1).

Epithelial stem cell proliferation and apoptosis are key to maintenance of normal intestinal homeostasis and epithelial membrane integrity. Given the histology differences seen between the diet groups, we next examined homeostatic responses including the endogenous nuclear protein Ki67, as a marker of cell proliferation through immunofluorescent staining. The MD had the highest Ki67 positive cells compared to CO (**Figure 2A and 2D).** Dysfunctions in apoptosis have been implicated in the pathogenesis of IBD (Nunes, Bernardazzi and de Souza, 2014) because without cell clearance by apoptosis, secondary necrosis occurs leading to increased intestinal permeability and ultimately chronic inflammation (Opferman and Korsmeyer, 2003). We examined apoptosis using the terminal deoxynucleotidyl transferase dUTP nick end labeling (TUNEL) assay for apoptotic DNA fragmentation (**Figure 2B and 2E**). The MD and MF diets show more cells positive for apoptotic DNA fragmentation than the CO diet suggesting balanced intestinal homeostasis.

**Figure 2.**
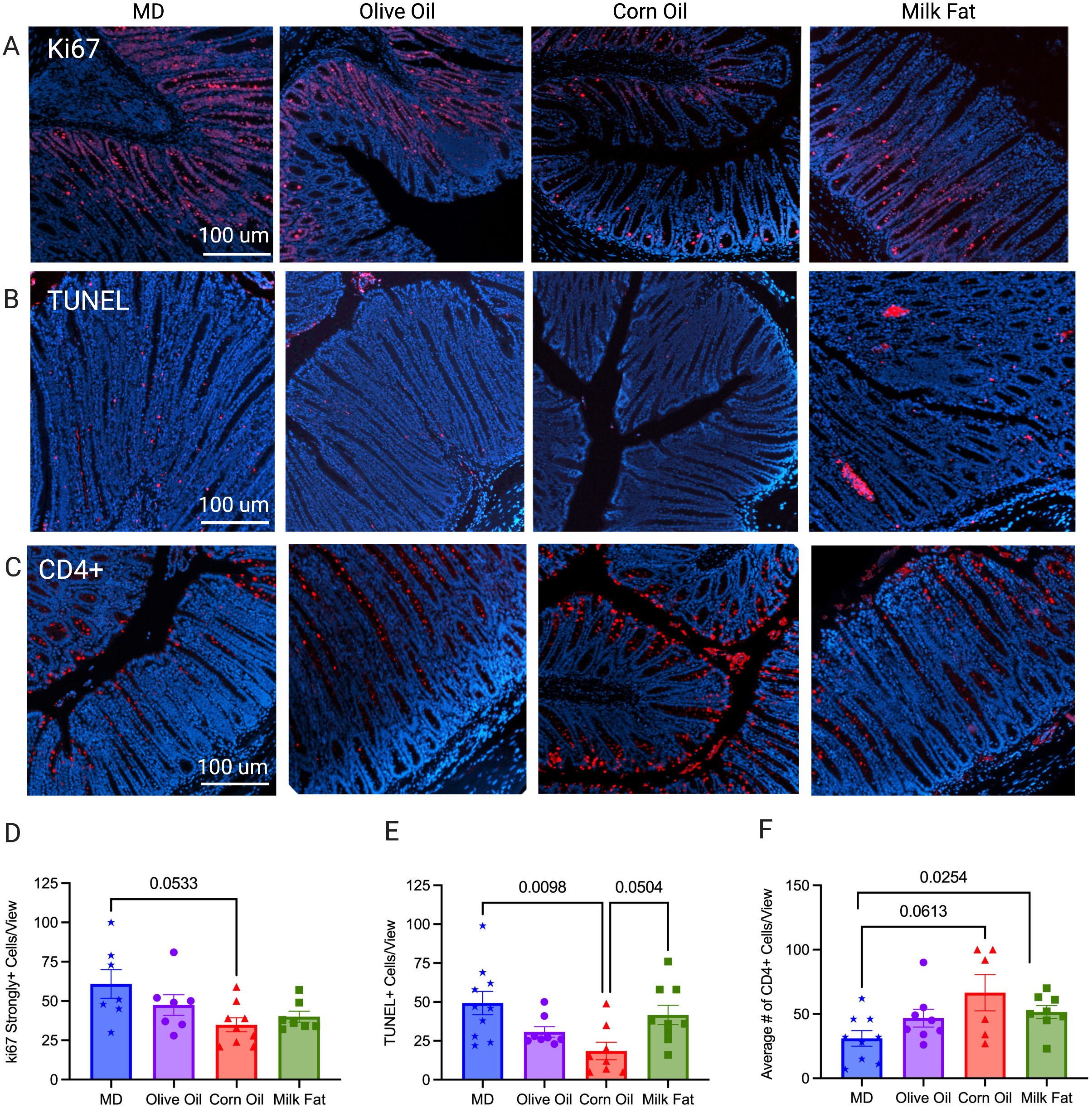
MD maintains tissue homeostasis through control of Ki67 and apoptosis. (A) Ki67 strongly positive cells seen as red florescence with DAPI used as a counterstain in the MD, olive oil, corn oil and milk fat diets, (D) quantification of the strongly positive Ki67 cells per view are shown. (B) Distal colon sections were stained for apoptotic cells using the TUNEL assay, strongly positive cells seen as red florescence with DAPI used as a counterstain and (E) the quantification number of apoptotic cells per view. (C) Immunostaining for CD4+ T-cells seen as red with DAPI used as counterstain and (F) the average number of CD4+ T-cells per view. *n*=8-10 mice/group. (scale bars: 100 μm, magnification, 100x). Statistical significance was determined by Kruskal-Wallis with Dunn’s post-hoc analysis, means ± SEM. MD: Mediterranean diet; TUNEL: terminal deoxynucleotidyl transferase dUTP nick end labeling; SEM: standard error of mean. *p*<0.05. (See also Figure S2).

Neutrophil infiltration was assessed through immunofluorescent staining of myeloperoxidase positive (MPO+) cells, as an over exuberant neutrophil response is known to cause epithelial damage in IBD (Williams and Parkos, 2007). The MF and OO fed diets had the highest number of MPO+ cells, with the MD showing significantly less infiltration than the MF diet (**Figure S2**). Unexpectedly, the CO fed diet had weak staining for MPO+ cell infiltration in comparison to the other diet groups despite showing more ulcerations and a more severe colitis which suggest other intestinal responses beside neutrophils are driving colonic damage although we did not rule out netosis. Immunostaining for T lymphocytes confirmed a greater influx of CD4+ T lymphocytes into the intestinal mucosa of the CO and MF fed diets in comparison to the MD (**Figure 2C and 2F**). CD4+ T cells are enriched in lesional tissue and are key initiators of disease progression in colitis (Imam *et al.*, 2018). The increased damage seen in the CO diet may be a result of suppression of important homeostatic immune responses (proliferation and apoptosis), and increased infiltration of CD4+ T lymphocytes. Overall, the fat blend contained in the MD protects against the development of severe and damaging colitis due to functional homeostatic balance.

### MD decreases colonic cytokines that drive colitis

The initiation, progression and resolution of inflammation observed in colitis are a result of cytokines which control multiple aspects of the inflammatory response (Neurath, 2014). An imbalance between pro-inflammatory and anti-inflammatory cytokines results in disease progression and tissue damage and limits the resolution of inflammation. To determine how the MD fat blend protects against the development of severe colitis, we examined the cytokine mRNA gene expression in the distal colon. In accordance with the clinical markers of disease and the histological analysis, we saw a significant decrease in the expression of mRNA inflammatory cytokines RELM-ß and IL-6 in the MD compared to the CO diet (**Figure 3A and 3B**). RELM-ß and IL-6 are known drivers of colitis in the Muc2^−/−^ model (Morampudi *et al.*, 2016). Despite the increased DAI and histological damage seen in the MF diet, the MF diet uniquely showed reduced expression of RELM-ß, yet increased expression of the antimicrobial peptide REG3-γ compared to the CO diet (**Figure 3A and 3C)**. No differences were seen between TNF-α, FOXP3, TGF-β1, Ebi3 or IL-22 colonic mRNA expression amongst the diet groups (**Table S3**). Taken together, the MD reduced the expression of the colonic mRNA pro-inflammatory cytokines that drive colitis in the Muc2^−/−^ model with a subsequent increase in localized colonic protective immune responses like REG3-γ.

**Figure 3.**
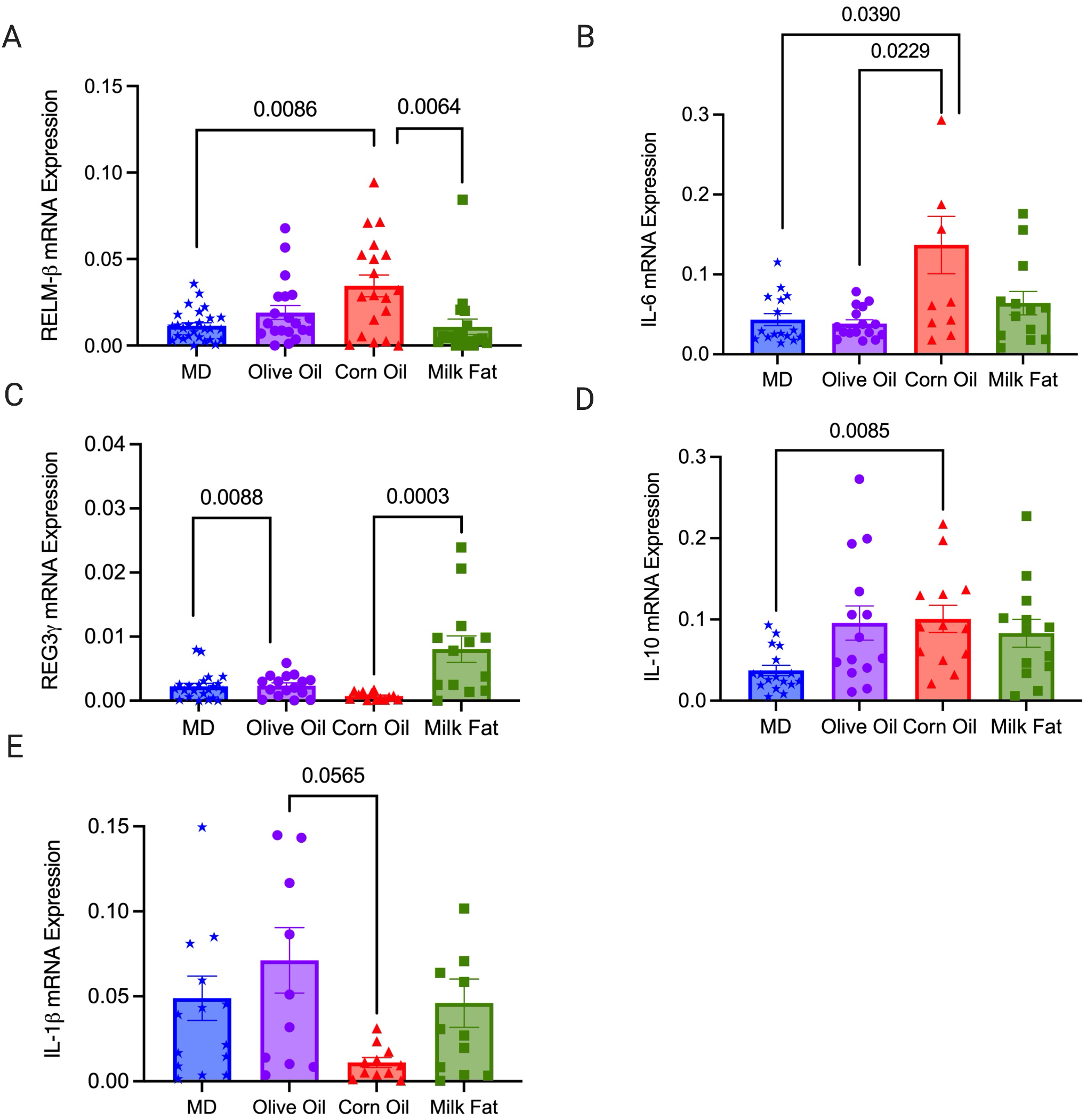
MD decreases colonic mRNA expression of pro-inflammatory cytokines RELM-ß and IL-6. Relative colonic mRNA expression (A) RELM-β, (B) IL-6 (C) REG3-γ, (D) IL-10 and (E) IL-1β. *n*=20-26 mice/group. Statistical significance was determined by Kruskal-Wallis with Dunn’s post-hoc analysis, means ± SEM. MD: Mediterranean diet; standard error of mean. *p*<0.05. (See also Figure S3).

### MD stimulates Th 22 cells important in maintaining epithelial homeostasis

Given that various types of fat have differential effects on the development of localized inflammation, we hypothesized that there were alterations in the composition of both innate and adaptive cells in the colon. We used immunophenotyping to determine the relative abundance of different immune cells found in the lamina propria of the colon. There were no significant differences between the diets in the proportion of inflammatory monocytes and neutrophils (**Table S4**), however the MF diet did show the largest absolute number of these cells, which correlates with the increased ulcers and abscesses seen in the histology. There were significant increases in macrophages in the MF diet compared to the MD and OO diets (**Figure 4A**), with significantly lower numbers of eosinophils in the MD in comparison to OO and MF diets. (**Figure 4B**). Eosinophils have been found in the inflamed tissue of colitis patients, leading to increase diarrhea, inflammation, and tissue destruction (Al-Haddad and Riddell, 2005). The cytokine IL-5 contributes to the detrimental effects of eosinophils, as eosinophils have specific receptors for IL-5 (Neurath, 2014). We see reduced eosinophils and serum IL-5 (**Figure 5D**) in the MD in comparison to the mice fed CO, further demonstrating the protective immunomodulatory effects of a fat blend (MD), favoring a milder colitis phenotype.

**Figure 4.**
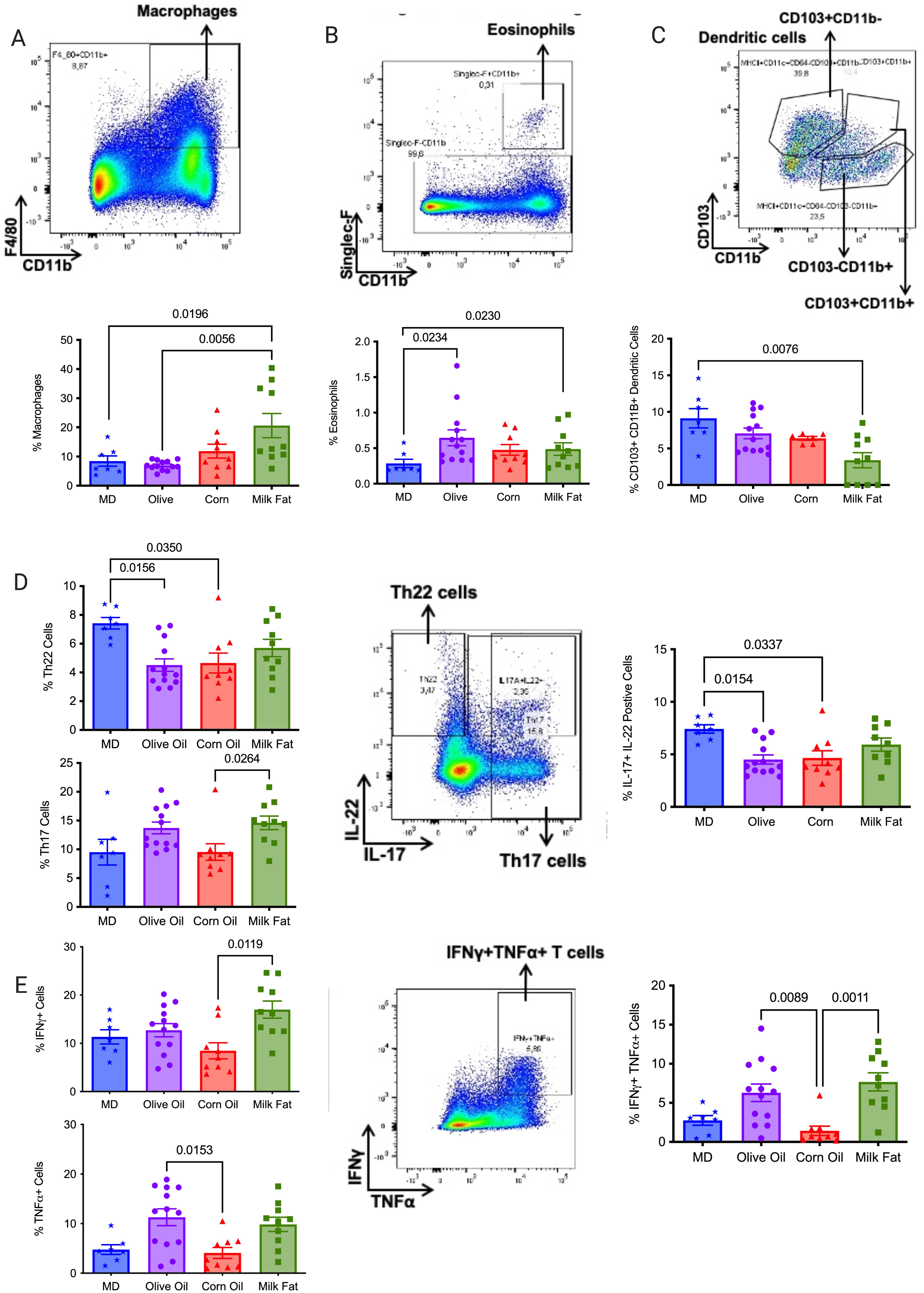
MD stimulates tolerogenic CD103^+^ dendritic cells and Th 22 cells that are vital in maintaining epithelial homeostasis. Flow cytometry analysis of the populations of intestinal lamina propria cells and their cytokines produced in the distal colon. The proportion of (A) macrophages (F4/80+CD11b+), (B) eosinophils (Siglec-F CD11B+), (C) dendritic cells (CD103+CD11b+), (D) Th22, Th17 and Il-17+IL22+ cells, (E) IFNγ+ cells, TNFα cells and IFNγ+TNFα+ cells. *n*=7-10 mice/group Statistical significance was determined by Kruskal-Wallis with Dunn’s post-hoc analysis, means ± SEM. MD: Mediterranean diet SEM: standard error of mean. *p*<0.05. (See also Figure S4).

**Figure 5.**
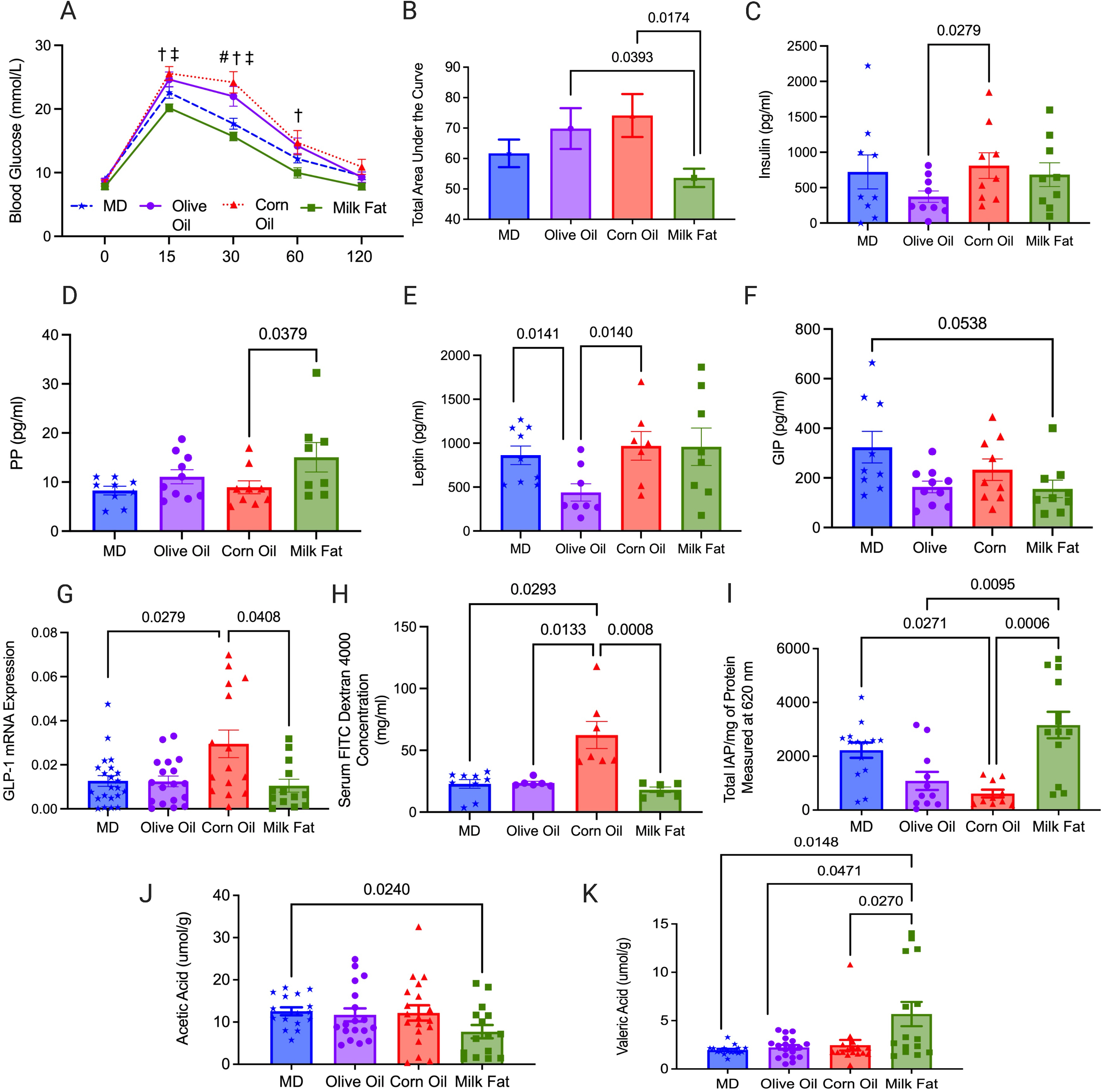
MD stimulates key cytokines G-CSF and MCP-1 necessary for a balanced immune response. Serum samples were collected at the end of the study. (A) G-CSF, (B) MCP-1, (C) IL-10, (D) IL-5, (E) IL-17 and (F) IFN-γ. The level of cytokine production in the serum were measured using by addressable laser bead immunoassay. *n*=8 mice/group. Statistical significance was determined by Kruskal-Wallis with Dunn’s post-hoc analysis, means ± SEM. MD: Mediterranean diet; OO: olive oil; CO: corn oil and MF: milk fat; SEM: standard error of mean. *p*<0.05. (See also Figure S5).

Dendritic cells (DC), in particular tolerogenic CD103^+^ DCs are crucial for intestinal homeostasis, as they prevent an aberrant immune response (Stagg, 2018). MD resulted in a significant increase in the tolerogenic CD103+CD11b+ DCs versus the MF group (**Figure 4C**). We next examined the various immune cell sub-types to further elucidate how type of fat impacts the adaptive inflammatory responses. Th17 and Th22 cells, along with serum IL-17 and MCP-1 play a protective role in IBD promoting barrier function through epithelial cell regeneration, host protection through immune cell recruitment and maintaining intestinal homeostasis (Gálvez, 2014; Lindemans *et al.*, 2015). Within the T helper (Th) cell populations Th22 cells and IL-17+IL-22+ cells were significantly increased in the MD versus the OO and CO fed diets (**Figure 4D)**. Th22 cells and IL-17+IL-22+ cells promote barrier repair and epithelial homeostasis (Lindemans *et al.*, 2015). Surprisingly, the CO diet had the most colonic damage and increased colonic CD4+ T cells, yet we observed a reduced proportion of Th17 and IFNγ+TNFα+ producing cells in the lamina propria (**Figure 4E**) with concomitant reductions in serum IL-17 and IFN-γ levels (**Figure 5E and 5F**). Further examination of the systemic cytokines indicate that the CO fed mice had reduced serum production of granulocyte colony stimulating factor (G-CSF) and monocyte chemoattractant protein-1 (MCP-1) when compared to the MD fed mice (**Figure 5A and 5B**). G-CSF is a key regulator of neutrophil differentiation and enhances the bactericidal function of neutrophils, whereas MCP-1 is a necessary component of the inflammatory response required for tissue protection, remodeling, and healthy expansion (Cranford *et al.*, 2016). Mice lacking G-CSF have been found to more susceptible to DSS-induced colitis (Meshkibaf *et al.*, 2016). This would support the weak staining for MPO+ cell infiltration for neutrophils with immunofluorescent staining in the CO diet in comparison to the other diet groups (**Figure S2A)**. Colonic mRNA expression of IL-10 and serum IL-10 were significantly increased in the mice fed the CO diet (**Figure 3D and 5C)**. IL-10 is considered an essential anti-inflammatory cytokine crucial in maintenance of immune homeostasis in IBD, however, there is emerging evidence that IL-10 may play a previously underappreciated dual role, with its function highly dependent on timing of IL-10 production(Couper, Blount and Riley, 2008). Our data shows that IL-10 mRNA expression was significantly upregulated in the CO diet compared to MD. In this scenario, as we discovered severe intestinal damage and more severe colitis in the CO diet, this could indicate a dysfunctional compensatory effect of IL-10 in this diet group given the associated damage in the colon. No differences in additional chemokines or cytokines were noted (**Table S5**).

Overall, the MD fat blend was more protective against dysfunctional immune responses than any fat alone. Although the MD did experience inflammation, the inflammation was protective and induced tissue repair which is essential in Muc2^−/−^ mice lacking the mucus layer.

### Milk fat improves glucose homeostasis, intestinal permeability, and barrier function

The Muc2^−/−^ mice develop metabolic dysfunction (Ye *et al.*, 2021), so we conducted an intraperitoneal glucose tolerance test (IPGTT) to investigate how the MD fat blend impact glucose tolerance compared to the individual fat diets. Baseline fasting glucose levels were not different between the various diet groups (0 minutes in IPGTT, **Figure 6A**), however upon glucose challenge, the mice fed CO and OO diets had significantly higher glucose levels than the MD and MF diets at multiple time points (**Figure 6A**). These results were confirmed by calculation of the area under the curve (**Figure 6B**) which is indicative that dietary fats have differential effects on glucose clearance.

**Figure 6.**
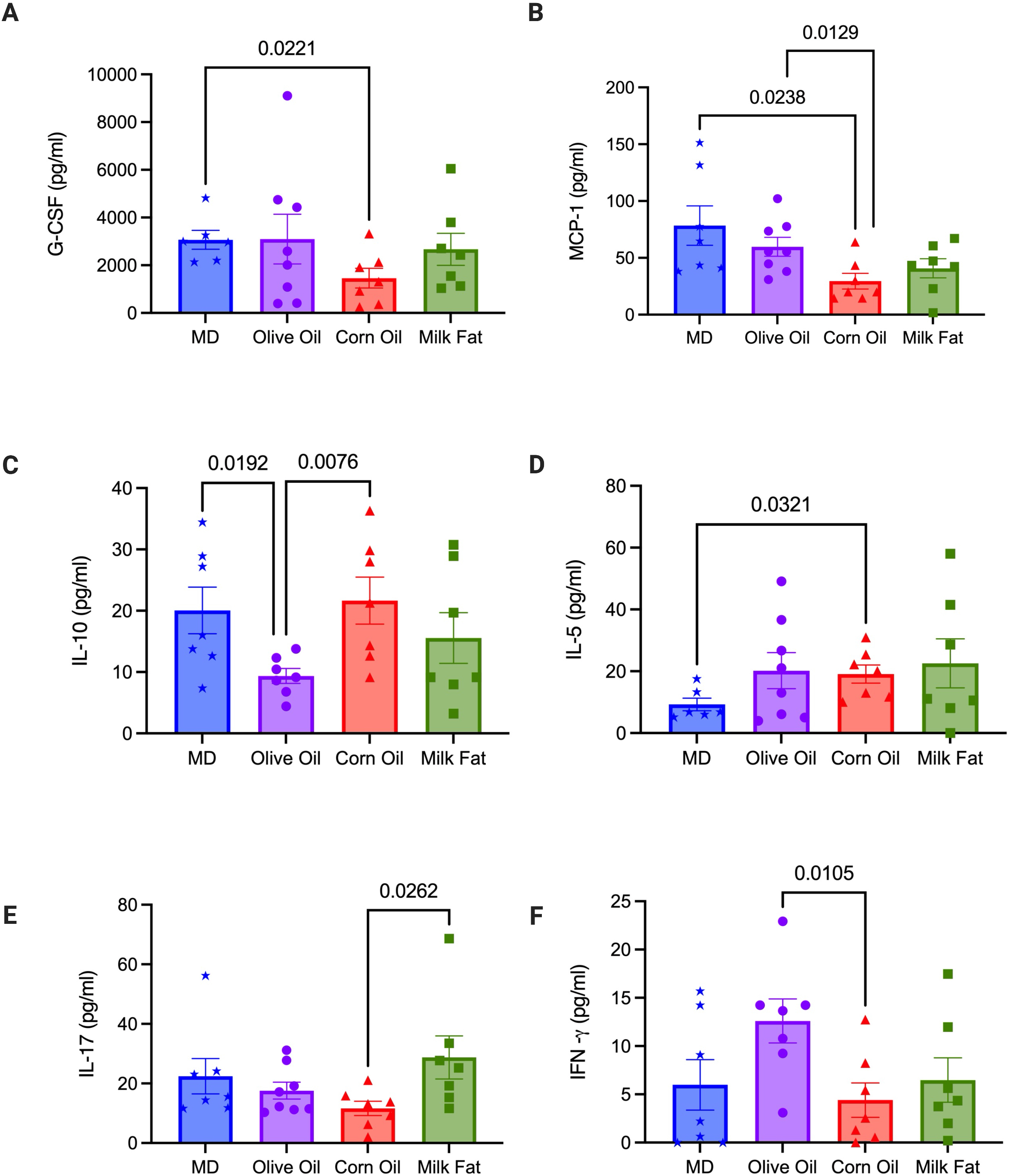
Saturated fat, derived from MF, improves glucose homeostasis, intestinal permeability, and barrier function. (A) Intraperitoneal glucose tolerance test (IPGTT) for all diet groups (*n*=12-19 mice/group) was performed at the end of the study, (B) along with corresponding area under the curve (AUC). Serum concentrations of hormones (C) insulin, (D) PP (E) leptin and (F) GIP (*n*=9-10/group). (G) mRNA expression of GLP-1. (H) *Fluorescein isothiocyanate*–*dextran* 4kDa (FITC) measured intestinal permeability (*n*=6/group). (I) The key mucosal defense factor intestinal alkaline phosphatase (IAP) measured in the colon (*n*=11-14 group). Short chain fatty acids (J) acetic acid and (K) valeric acid were measured in the cecum (*n*=15-19 mice/group). MD: Mediterranean diet; PP: pancreatic polypeptide; GLP-1: glucagon like peptide-1; GIP: gastric inhibitory peptide; SFA: saturated fatty acids; MUFA: monounsaturated fatty acids; n-6 PUFA; n-6 polyunsaturated fatty acids; n-3 PUFA: n-6 polyunsaturated fatty acids; SEM: standard error of mean. ^§^*p*<0.05 than milk fat; ^‡^*p*<0.05 than corn oil;#*p*<0.05 than MD; and ^†^*p*<0.05 than olive oil. (See also Figure S6).

Circulating serum metabolic hormone levels revealed the OO diet had significantly lower serum insulin levels than the CO diet (**Figure 6C)** and pancreatic polypeptide (PP) was significantly increased in the MF diet versus the CO diet (**Figure 6D)**. Leptin was significantly lower in the OO diet compared to the MD and CO diet (**Figure 6E)**. The MD had significantly higher serum levels of glucose inhibitory peptide (GIP) in comparison to the MF diet (**Figure 6F**). Colonic mRNA gene expression of glucagon like peptide 1 (GLP-1) demonstrated a significant increase in expression in the CO diet when compared to the MD and MF diets (**Figure 6G)**, however no changes were observed in serum GLP-1 which is likely a compensatory signal to promote increased synthesis of serum GLP-1. No differences were seen in the serum levels of glucagon, amylin, peptide YY and C-peptide, ghrelin, resistin or GLP-1 between the diet groups (**Table S6**). In summary, serum insulin levels between the MD, MF and CO diet are similar, yet result in vastly different glucose responses. Despite the consistent levels of insulin between the MD, MF and CO diets, the CO diet requires more insulin to maintain euglycemia than the other groups, therefore it plausible that a CO diet induced insulin resistance. To further support this, MCP-1 deficiency in the CO diet further contributed to the metabolic perturbations which is consistent with previously reported findings (Cranford *et al.*, 2016), where MCP-1 depletion has been linked to high fat diet pathologies, including metabolic dysfunction and fibrotic adipose tissue. In contrast to CO, in the OO diet, we see impaired glucose homeostasis due to reduced serum insulin and leptin secretion. Glucose homeostasis is closely regulated by both insulin and leptin, with decreased leptin potentially contributing to reduced insulin sensitivity in the OO diet (Paz-Filho *et al.*, 2012).

Mounting preclinical evidence in colitis supports the contribution of intestinal permeability (IP) to increased systemic inflammation in metabolic diseases as a result of alterations to the gut microbiome (Madsen *et al.*, 1999). We sought to determine if the alterations in immune responses and metabolism were a result of changes in IP. Through the FITC dextran assay, we demonstrate that the MD, OO and MF diets demonstrated similar IP, whereas the CO diet had increased IP (**Figure 6H**). We next examined intestinal alkaline phosphatase (IAP), as IAP plays a major role in the maintenance of intestinal homeostasis and protection with reduced expression associated with worsened mucosal inflammation and metabolic syndrome (Lallès, 2010). We sought to determine if IAP enzymatic expression would differ based on the type of fat provided in the diet. The MD and MF diets show increased IAP enzymatic expression in comparison to the CO and OO diets (**Figure 6I**) suggesting that saturated fat plays a key role in IAP enzyme expression.

Short chain fatty acids (SCFA) including acetic, propionic, and butyric acid have important immunomodulatory properties and promote gut homeostasis (Parada Venegas *et al.*, 2019), we examined the cecal production of SCFA. No significant differences were seen in total SCFA, butyrate or propionate (**Figure S6A to S6C**). However, the MF diet did show reduced production of acetic acid in comparison to the MD (**Figure 6J**) and increased production of valeric acid (**Figure 6K**) in comparison to all diet groups. Although further work is needed to elucidate the influence of valeric acid and acetic acid on intestinal inflammation, preliminary evidence suggests that valeric acid confers protective effects against radiation-induced colitis (Y. Li *et al.*, 2020), yet reduced levels of acetic acid could enhance susceptibility to colitis (Laffin *et al.*, 2019).

In summary, the results presented above indicate that type of fat influences glucose homeostasis, serum hormones, IP, SCFA and IAP expression differentially. Diets that contain saturated fat, such as the MD and MF diets demonstrate improvements in glucose metabolism, which is supported by IPGTT, serum hormones, increased expression of IAP and improved IP. However, diets rich in CO and OO contribute to impairments in metabolism as seen by alterations in glucose tolerance, reduced expression of IAP and the CO diet also showing impaired IP. Overall, the MD fat blend containing SFA is an important part of the beneficial effects of the diet metabolism in a colitis susceptible host.

### The MD promotes a shift away from a colitis-associated microbiome abundant in health promoting bacteria

The emerging evidence on the interconnectedness between the gut microbiome and host metabolism (Visconti *et al.*, 2019), as well as the hypothesized role of the microbiome in the etiology of IBD led us to investigate how the MD fat blend influenced the composition of the gut bacteriome in Muc2^−/−^ mice compared to the individual fat diets. Through 16rRNA gene sequencing of the V4-V5 region, we characterized the bacteriome of the colon post diet treatment. Thirty-eight samples were successfully sequenced (7 CO, 8 MD, 9 MF and 14 OO). No significant effect of cage, sex, or sequencing batch were detected by using a PERMANOVA test on the robust centered log-ratio transform (**Figure S7A)**. No differences in α-diversity were observed across diet groups using the metrics of observed number of unique ASVs (richness), Pielou’s evenness, Shannon diversity or Faith’s phylogenetic diversity (PD) (**Table S7B**). β-diversity distances calculated from a robust cantered-log ratio transform of the ASVs relative abundance table and tested using a PERMANOVA determined differences between diet groups (p=0.003, pseudo-F=3.6, permutations=999) (**Figure 7A**). A post-hoc pairwise test of the PERMANOVA results revealed that the CO group was statistically significant from the MD and OO, and appreciably different than MF, though this was not statistically significant (**Table S7C**). Differential abundance testing was guided using Songbird and the differential ranks visualized using Qurro (**Figure S7D**). The top ASVs associated with the MD were *Absiella innocuum*, *Alistipes sp.*, *Emergencia timonensis* and *Lactobacillus animalis*, whereas *Blautia massiliensis*, *Faecalibaculum rodentium, Akkermansia muciniphila, Anaerosporobacter mobilis* were associated the CO diet (**Figure 7B**). When the MD was compared to the MF diet, the top ASVs associated with the MD were *Alistipes sp., Lactobacillus animalis, Olsenella 001457795, Desulfovibrio fairfieldensis and Faecalibaculum rodentium* versus in the MF diet *Blautia massiliensis, Dorea faecis, Lawsonibacter sp.* and *Ruthenibacterium lactatiformans* and were most abundant (**Figure 7C**). The top ranked ASVs in the MD when compared to OO diet *Paramuribaculum intestinale*, *Odoribacter massiliensis*, Muribaculaceae and *Alistipes sp.* versus *Dorea faecis*, *Bacteroides massiliensis*, *Muribaculum sp003150235* and *Absiella innocuum* were most abundant in the OO diet (**Figure 7D**). Overall, there was a consistent association with Enterobacteriaceae and *Bacteroides massiliensis* in all the fat diets compared to the MD. The effect of the CO diet on different ASVs was most pronounced out of all the diet groups with OO being most similar. Together, these results confirm that various types of fat alter the composition of the gut microbiota.

**Figure 7.**
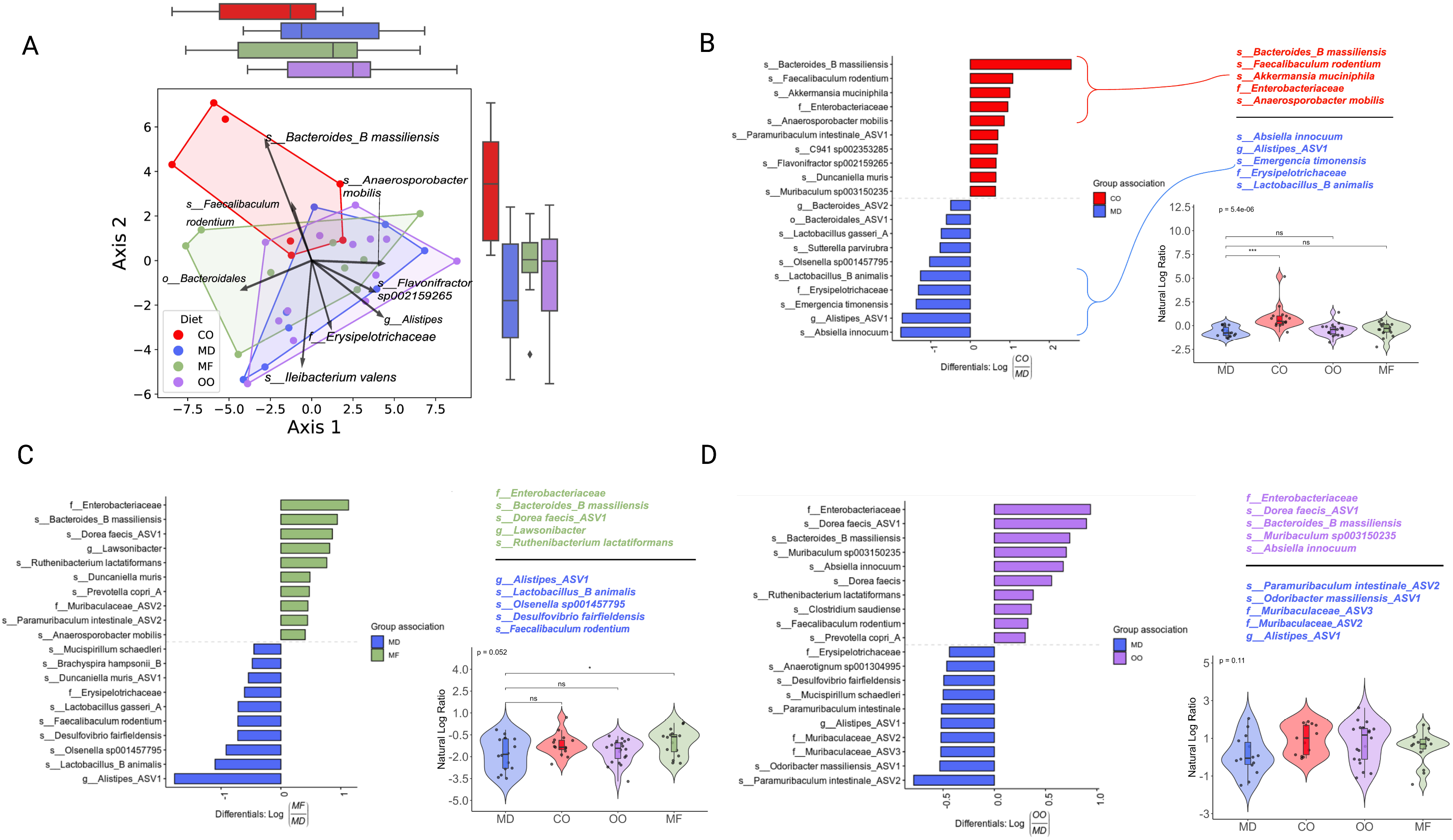
The MD promotes a shift away from a colitis-associated microbiome abundant in health promoting bacteria. Colon samples collected post-diet intervention and sequenced for 16S rRNA at the V4-V5 region. (A) Principal component analysis biplot of robust centred-log ratio transformed distances. Circles represent individual mice, vectors represent ASV loadings, and the diet groups are enclosed with convex hull polygons. The ridge boxplots represent the principal component values along the corresponding axis. An overall significant difference between groups was assessed using a PERMANOVA test (*p*=0.003, Pseudo F=3.6, permutations=999) and the assumptions of heterogeneous dispersion were checked using a PERMDISP test (*p*=0.769, Pseudo-F=.77, permutations=999). Pairwise post-hoc testing shows a significant difference between the CO and MD groups, as well as CO to OO groups. (B to D) Ranked plots of the inverse additive log-ratio transform (inverse ALR) differentials from Songbird’s multinomial regression analysis, which estimates the probability of an ASV being observed for a specific diet group. The top and bottom 10 ranked ASVs are displayed as their highest classified taxonomic level based on the GTDB reference database. A positive value indicates higher association with the numerator group, and a negative value indicates higher association with the denominator group, the MD is used as the reference group compared to CO, MF and OO diets (n=7-14 mice per group). (See also Figure S7). ASV: amplicon sequence variant; GTDB: gene taxonomy database; PERMANOVA: permutational multivariate analysis of variance; PERMDISP: permutational analysis of multivariate dispersions.

## Discussion

The role of diet in IBD is underappreciated, yet food is essential to nourishment, socialization, and well-being in this population. Evidence-based diet recommendations are needed to determine which dietary patterns IBD patients can tolerate in their everyday lives. Despite the increasing evidence of the influence of dietary components and its impact on the disease activity, inflammation and the microbiome, large gaps still exist. This mechanistic study provides compelling evidence that type of fat, not total calories derived from fat, play a central role in shaping the microbial ecology, inflammation, and disease activity in a murine model (Muc2^−/−^) that develops spontaneous colitis with the MD fat blend protecting against severe colitis.

In this study, we show that MD, created by blending high levels of MUFA (from OO), SFA (from MF) combined with some n-3 PUFA (from fish oil) with low n-6 PUFA (from CO) creates a protective fat combination against colitis. Mice fed this MD fat blend were protected from developing severe colitis demonstrated by reduced disease activity, histological damage in the distal colon, including less ulcers and abscesses compared to the other diet groups. In contrast, mice fed a diet composed of CO or MF develop severe colitis with increased mucosal damage. Although the MD is rich in OO, OO alone resulted in the Muc2^−/−^ mice developing colonic damage, albeit less severe than that seen in the CO or MF diets which both resulted in severe colitis. This data exemplifies that the MD fat blend, not a single type of fat, is important in its ability to completely protect against colitis.

Inflammation was investigated both locally and systemically with each type of fat demonstrating different immunological responses. Unique to the MD, we see a significant increase in the proportion of Th22 cells and tolerogenic CD103+CD11b+ dendritic cells, as well as immune regulating signals like serum G-CSF, which are critical in mediating tolerance to antigens, limiting reactivity to the gut microbiota and are required for restitution (Meshkibaf *et al.*, 2016; Imam *et al.*, 2018; Stagg, 2018). Similar to the MD, the MF diet showed compensatory protective responses, such as increased expression of the key mucosal defense factor IAP and antimicrobial lectin RegIII-□ that promotes restitution important in preventing the cycle of chronic inflammation (Ramasamy *et al.*, 2011; Loonen *et al.*, 2014). These data suggests that the inflammatory response, as seen in the MD, is counterbalanced by an immunosuppressive response, which limits colonic damage. When MF is provided within a blend of fats, as seen in the MD, the protective inflammatory responses of the MF diet are conferred, limiting mucosal damage and ultimately colitis severity.

Dietary fat composition has been implicated in the development of insulin resistance, including the development of type 2 diabetes, with each type of fat having vastly different effects on insulin resistance and metabolic control (Ikemoto *et al.*, 1996). Curiously, metabolic abnormalities are increasingly being identified in both human and animal models of colitis (Verdugo-Meza *et al.*, 2020). We demonstrate that in Muc2^−/−^ mice that OO and CO diets promote impairments in glucose tolerance, barrier function and alterations to hormone levels leading to metabolic alterations. Both the MD and MF diet, composed of SFA, demonstrate improved glucose tolerance and barrier function. In line with the unique immunological effects of each type of fat, these results demonstrate that MF provided as component of a fat blend (MD) plays a key role in maintaining glucose homeostasis and gut barrier function. The potential mechanisms could be through increased expression of the key mucosal defense factor IAP (Lallès, 2010) and upregulation of the antimicrobial lectin RegIII-□ (Morampudi *et al.*, 2016), as well as increases in key hormones like PP known to enhance insulin sensitivity (Rabiee *et al.*, 2011).

It is well documented that alterations in the microbiota are linked to IBD, metabolic diseases, as well as other inflammatory conditions (Verdugo-Meza *et al.*, 2020). Multiple research groups have established the effect of a high fat diet (i.e., the total calories derived from fat) on the microbiome (Teixeira *et al.*, 2011; Lee *et al.*, 2017; Li *et al.*, 2019), however knowledge on the influence of specific types of fat on gut microbial ecology is lacking. Our analyses demonstrate that not only type of fat alters the composition of the gut microbiota but that the composition of the fat found in the MD contributes to beneficial changes in the microbiota and subsequent protective health effects. Top ranked taxa associated with MD consisted of ASVs classified as *Absiella innocuum*, *Lactobacillus animalis, Emergencia timonensis* and *Alistipes sp.. L. animalis* is a well-known beneficial microbe recognized to contribute to immune modulation, epithelial adherence, enhancement of gut barrier function and anti-tumorigenic potential (Hibberd *et al.*, 2017). Moreover, *Alistipes* spp. have been associated with less severe colitis (Butera *et al.*, 2018). Uniquely, the MD and the OO diets had a higher ratio of ASVs from the family of the health-promoting microbe Muribaculaceae which has been associated with extreme longevity in rodents (Sibai *et al.*, 2020). Additionally, accumulating evidence that the presence of Muribaculaceae in the gut is negatively associated with chronic diseases of the industrialized world (Sonnenburg and Sonnenburg, 2019) and its disappearance in the microbiota is associated with the increased incidence and prevalence of chronic diseases such as diabetes (Livanos et al., 2016). In contrast, the top ranked ASVs in the CO, MF and OO diets were *B. massiliensis*, with the CO diet also containing *A. muciniphila* both of which have been associated with colitis and colorectal cancer in humans and animal models (Berry *et al.*, 2012; Baxter *et al.*, 2014; Feng *et al.*, 2015). *A. muciniphila* is generally thought of as a health-promoting microbe (de Vos, 2017), however with alterations to the protective mucous layer, this microbe has pathobiont characteristics (Ganesh *et al.*, 2013). As *A. muciniphila* are known mucin-degraders, combined with a disrupted mucus layer as seen in the Muc2^−/−^ mouse model, it is likely pathobiont that is able penetrate the mucosal layer resulting in mucosal damage (Crost *et al.*, 2013). The higher ratio of Enterobacteriaceae in the CO, MF and OO diets and *Dorea faecis* observed in the MF and OO diets may have also contributed to colitis, as others have seen an enrichment in these microbes in spontaneous colitis as they are known to promote chronic intestinal inflammation (Bloom *et al.*, 2011; Berry *et al.*, 2015). The finding that there were increased ratios of ASVs of both mucin-degrading and pro-neoplastic bacteria, is clinically important in IBD, as the risk of colorectal cancer increases when there is exposure to prolonged chronic intestinal inflammation (Stidham and Higgins, 2018). The precise role of type of fat and its role in colitis and the gut microbiome, should be confirmed in human studies.

In summary, this study shows that a diet with a fat-blend, as such seen in a Mediterranean diet significantly reduces disease activity, inflammation-related biomarkers and improves metabolic parameters in the Muc2^−/−^ mouse model. In addition, we show that each type of fat differentially impacts the development of colitis and that it is not necessarily the total fat content of the diet that aggravates colitis. Although, the observed effects of a MD need to be confirmed through interventional studies in individuals living with IBD, it would be prudent to have patients focus on the types of fatty acids and their subsequent food sources versus restriction of total fat in the diet. The recommendation to reduce fat intake in the diet, in particular SFA, are not aligned with current evidence (Astrup *et al.*, 2020) and as such could reduce intake of important nutrient-dense foods in a population that is at risk for nutrition deficiencies. The MD is a healthful diet that has the potential to maintain an appropriate immune response while mitigating the damaging effects of chronic inflammation and begs future research to confirm these results in humans.

## Acknowledgments

NH was funded by a Canadian Institutes of Health Research - Frederick Banting and Charles Best Canada Graduate Doctoral Award and a Canadian Association of Gastroenterology PhD Studentship Award. This study was supported by a Crohn’s and Colitis Canada Grant-in-Aid to DLG and an NSERC Grant-in-Aid to SG. All figures created in Biorender.com.

## Author Contributions

N.H., and D.L.G., conceived and designed the research, N.H., J.Y., J.A.B, A.A.M, M.Y., B.W.B., and M.E. executed the experiments. N.H. with input from D.L.G analysed the data with statistical advice and design input from M.E.. N.H. wrote the original draft of the manuscript and D.L.G. reviewed and edited the manuscript. D.L.G. supervised, provided resources and funding for this project. S. Ghosh and S. Gruenheid provided resources and expertise.

---

**Table 1. Nutrient composition of experimental diets.** Four diets were examined in this study – Mediterranean Diet, Olive Oil, Corn Oil and Anhydrous Milk Fat. All diets were isocaloric, iso-nitrogenous with similar protein, carbohydrate, vitamin and mineral profiles. Lipid content was altered to reflect the various fatty acid profiles.

## Materials and Methods

### Animals Models

Mucin 2 deficient (Muc2^−/−^) mice with a C57BL/6 wild-type background were originally obtained from the Vancouver Gastrointestinal Disease Research Program (B. Vallance lab, Vancouver, BC, Canada) and bred at UBC Okanagan and backcrossed with Charles River C57BL/6 mice. Mice were sex-matched and housed in sterilized, filter-topped cages, handled in a biological safety cabinet in a specific pathogen free environment with sentinels used to test for common pathogens. All mice had free access to acidified tap water (pH of ~2.3 via the addition of HCl) and fed irradiated food in a temperature-controlled room (22°C) with a controlled reverse lighting cycle (12-hour dark/light cycle). The animal experiment was approved by the University of British Columbia’s Animal Care Committee (Protocol No: A15-0240), in accordance with the Canadian Council on Animal Care Standards.

### Experimental Set-up

Animals were weaned at the age of 19 days, pooled into cages (n=3-4 mice/cage) and randomly divided into four diet groups which only differed by fat composition (20.2% w/w fat derived from corn oil (TD. 120022), olive oil (TD. 130128), anhydrous milk fat (TD. 120021) or a Mediterranean Diet like fat blend (TD. 170674) - Teklad Envigo, Madison, WI) according to sex with equal numbers of males and females included in the study. The fat blend was referred to as a Mediterranean Diet (MD) with the fatty acid profile designed to reflect the MD consumed by humans in Mediterranean regions (40.8% calories from fat with 7.8% SFA, 27.7% MUFA and 4.5% total PUFA (0.8% derived from n-3 PUFA) (Serra-Majem *et al.*, 2009). To reduce cage effects, the mice of the same sex were pooled into cages (*n =* 3–4 mice/cage). To mitigate husbandry effects, paired breeding (1 male and 1 female) was used, with the male staying with females and pups after birth. Upon weaning, pups were equally divided into cages, mixed with mice weaned from another litter, so in each cage, there was a mix of mice from different breeder pairs. Environmental cage effects were mitigated by monitoring well-being and environmental conditions (e.g., light, temperature, relative humidity) daily without disturbing the cage. Fight wounds, which are indicative of aggressive behavior, were not observed. Each cage had 67.6 inches of floor space, covered in woodchips, cotton nesting, and Nestlets for cage enrichment. Once per week, at the same time of day, body weight and clinical scoring was completed. To evaluate the clinical disease activity of colitis, body weight, stool consistency, rectal bleeding was recorded, and a disease activity score was calculated as previously described (Cooper *et al.*, 1993). At day 84, the mice were euthanized, and the tissues excised (colon, cecum, liver), flash frozen in liquid nitrogen, immersed in 10% formalin (Fisher) or RNAlater (Qiagen). Flash frozen tissues and those stored in RNAlater were stored at −80°C until further analysis.

### Histopathological Scoring

Colon tissues were embedded in paraffin and stained with hematoxylin and eosin by Wax-it Histology Services Inc., (Vancouver, BC, Canada). Coded samples blinded by the scorers were evaluated and scored by at least two people. The scoring of colonic inflammation was quantified using a combination of approaches as previously described (Rath *et al.*, 1996; Bergstrom *et al.*, 2010).

### Immunofluorescence

Paraffin-embedded colon tissue sections were deparaffinized in xylene and progressively rehydrated in decreasing concentrations of ethanol (100, 90, 80 and 70% for 3 min each), and finally incubated in de-ionized water for 3 min. The Ag retrieval process was performed by incubating the slides a 1 mg/mL trypsin (Sigma) for 30 min followed by a 5% BSA blocking solution. The sections were then incubated for 2 hours at room temperature using: rabbit polyclonal antibody-1 for myeloperoxidase (Invitrogen) to examine neutrophils; rat monoclonal antibody for F4/80 (CedarLane) to examine macrophages and rabbit monoclonal antibody for Ki-67 (CedarLane) for cellular proliferation. Secondary antibodies used include goat anti-rabbit IgG AlexaFluor-conjugated 594-red antibody (Invitrogen) or goat anti-rabbit IgG 488-conjugated antibody (Invitrogen). Tissue sections were mounted using fluoroshield with DAPI (Sigma) and viewed on an Olympus IX81 fluorescent microscope. For inflammatory cell counts, positive cells were quantified by two blinded observers under fluorescence using Olympus cellSens Software.

### TUNEL Assay

Apoptotic DNA fragmentation was examined using 30064 CF™ 594 Dye TUNEL Assay Apoptosis Detection Kit (Biotium, Cedarlane) according to the manufacturer’s protocol. Briefly, paraffin-fixed colon tissues were deparaffinized according to standard protocols. Cells were permeabilized in 20 ug/ml proteinase K in PBS at 37°C for 30 minutes, incubated for 5 minutes with 100 ul of TUNEL Equilibration Buffer, followed by TdT enzyme with TUNEL Reaction Buffer and labeled with fluorescein 594-dUTP using terminal deoxynucleotidyl transferase. The localized red fluorescence of the apoptotic cells was absorbed using Olympus cellSens Software.

### RNA extraction and Quantitative Real-Time PCR (qPCR)

Upon excision, colon and liver tissues were placed in RNAlater (Qiagen) and stored in the −80□ freezer until extraction. RNA was extracted from the tissues using the Qiagen RNeasy Fibrous Tissue Mini kit (Qiagen) according to the manufacturer’s instructions. Total RNA was quantified using a NanoDrop 2000c Spectrophotometer (Thermo Scientific) and cDNA synthesized with iScript cDNA Synthesis Kit (Bio-Rad). Quantification of cDNA was performed on a Bio-Rad CFX Manager 2.0 machine using Sso Fast Eva Green Supermix (Bio-Rad). All primers were synthesized by Integrated DNA Technology, Canada (**Table 2**). Gene expression was normalized to TBP mRNA level and calculated as ΔCt = 2(^CtTBP^ ^mRNA-CTgene^ ^of^ ^interest^ ^mRNA^).

### Serum Analyses

Blood was collected from the mice via cardiac puncture, serum removed and stored at −80°C. Protease inhibitor was added to sera (Protease Inhibitor Cocktail, VWR Life Science Amresco) and analyzed for metabolic hormones (Mouse Metabolic Array) and a panel of 31 chemokines/cytokines (Mouse Cytokine Array/Chemokine Array 31-Plex) by addressable laser bead immunoassay by Eve Technologies (evetechnologies.com; Calgary, AB, Canada).

### Flow Cytometry

#### Isolation of intestinal lamina propria cells

Intestinal lamina propria cells from mice were isolated using a modified version of a previously described protocol (Weigmann *et al.*, 2007) In brief, colons were removed and placed in cold calcium- and magnesium-free Hanks balanced salt solution (HBSS; Gibco) supplemented with 2% heat-inactivated FCS and 15 mM HEPES (Gibco). Intestines were cut open longitudinally, washed thoroughly, cut into 2 cm pieces, and incubated with shaking in EDTA buffer (HBSS supplemented with 2% FCS, 15 mM HEPES, and 5 mM EDTA) for 60 minutes at 37°C to remove epithelial cells. After removing the supernatant, tissue pieces were incubated in RPMI-1640 (Sigma) supplemented with 10% FCS, 15 mM HEPES, 100 μg/ml DNase I (Roche) and 200 μg/ml collagenase type IV (Sigma) for 40 minutes at 37°C. Cell suspension was filtered through a 70 μm cell strainer (Sigma), washed, and resuspended in FACS buffer (1X PBS supplemented with 2% FBS and 0.5M NA2EDTA) before proceeding with antibody staining.

For flow cytometry, the cells were incubated with 1 mg/ml rat anti- mouse CD16/CD32 Ab (Fc-block; clone 2.4G2) for 15 min at 4°C and then washed with cold FACS buffer. Fluorochrome-labeled extracellular antibodies were added in a total volume of 100 μl to 1×10^6^ cells, mixed thoroughly, and incubated for 25 minutes at 4°C. Extracellular antibodies used in this study are listed below in different panels and were used at a dilution of 1:200. Following extracellular staining, cells were washed with PBS and resuspended in viability dye (Life Technologies) for 20 minutes at 4°C. Intracellular staining for cytokines IFNγ, IL-17, IL-22 and TNF-α was performed using eBioscience™ Intracellular Fixation & Permeabilization Buffer Set. Prior to staining, cells were stimulated with Cell Stimulation Cocktail (plus protein transport inhibitors) (eBioscience) for 3 hours at 37°C. FoxP3 and RORgT staining was performed using eBioscience™ Foxp3 / Transcription Factor Staining Buffer Set and antibodies were used at a dilution of 1:100. A FACSCanto II (BD Biosciences) was used for sample analysis and flow cytometric analysis was performed using FlowJo software (TreeStar, Inc).

### Short-chain fatty acids (SCFAs) analysis

Direct-injection gas chromatography was used to quantify SCFAs acetic, propionic, butyric and valeric acid from cecal samples collected from Muc2^−/−^ mice (Schwarz *et al.*, 1996). Briefly, cecal samples were homogenized in isopropyl alcohol, containing 2-ethyl butyric acid at 0.01% v/v used as an internal standard and then centrifuged and the supernatant removed. The supernatant was injected into a Trace 1300 Gas Chromatograph, equipped with flame-ionization detector, with AI1310 autosampler (Thermo Scientific, Walkham, MA, USA) in splitless mode. A fused-silica FAMEWAX (Restekas, Bellefonte, PA, USA) column 30◻m × 0.32◻mm i.d. coated with 0.25◻μm film thickness was used. Helium was supplied as the carrier gas at a flow rate of 1.8◻ml/min. The initial oven temperature was 80°C, maintained for 5◻min, raised to 90◻°Cat 5◻°C/min, then increased to 105°C at 0.9◻°C/min, and finally increased to 240◻°C at 20◻°C/min and held for 5◻min. The temperature of the flame-ionization detector and the injection port was 240◻°C and 230◻°C, respectively. The flow rates of hydrogen, air and nitrogen as makeup gas were 30, 300 and 20◻ml/◻min, respectively. Data were analyzed with Chromeleon 7 software (Bannockburn, IL, USA). Fine separation of SCFA was confirmed by the complete separation of the volatile-free acid mix (Sigma, Oakville, ON, Canada). Data are presented as absolute values (g of SCFA per g of feces).

### FITC-dextran assessment of intestinal permeability

Mice were gavaged with FITC-dextran (molecular mass, 4 kDa; FD4; Sigma-Aldrich) at a concentration 80 mg/100 g body weight. Four hours after gavage, blood was collected by cardiac puncture, placed in 3% acid-citrate dextrose, and centrifuged at 1000 xg for 12 minutes to remove serum. Fluorescence of FITC-dextran in serum was diluted to 1:10 with PBS and measured on a Promega GloMax Multi Detection System (Promega) at 490 nm excitation and 520 nm emission wavelengths. FITC-dextran concentration was determined from a standard curve generated by serial dilutions of FITC-dextran.

### Intestinal Alkaline Phosphatase Assay (IAP)

IAP was extracted from 25 mg colon tissues in 500 ul of RIPA buffer (50 mM Tris, pH 8.0, 1% Triton-X 100, 0.5% sodium deoxycholate, 0.1% sodium dodecyl sulfate, 150 mM sodium chloride) and protease inhibitor. The sample was homogenized at 30Hz for 2 minutes then centrifuged at 1610 ×g for 5 min, and the supernatant containing IAP was collected. The IAP assay was completed using the Alkaline Phosphatase Activity Fluorometric Assay Kit (BioVision) according to the manufacturer instructions. Briefly, each 10 ul of each sample was treated with 100 ul of ALP Assay Buffer and 20 ul of 0.5 mM Methylumbelliferyl phosphate disodium salt (MUP) and incubated at 25°C for 30 minutes. Next, 20 ul of an inhibitor (aqueous K2HPO4) was added to each well. In addition, a standard curve was created using stock solution of 1 mg/mL BSA (Sigma) in triplicates with multiple concentrations. Florescence intensity was measured using the Promega GloMax Multi Detection System (Promega) at 360 nm excitation and 440 nm emission. Total relative protein concentration was quantified using the Bio-Rad Protein Assay (Bio-Rad, Ontario, Canada). IAP values are expressed as units of IAP/mg of protein.

### Glucose Tolerance Test (IPGTT)

At nine weeks, an intraperitoneal glucose tolerance testing (IPGTT) was completed, using standard protocols from the D.I.A.B.E.T.E.S. Centre (UBC-Okanagan, Kelowna, BC, Canada). Briefly, a dose of 1g/kg body weight glucose (20% wt/vol glucose solution) was administered via intraperitoneal injection. Blood sampling and glucose testing was performed at 0, 15, 30, 60, 90 and 120-minutes following glucose injection.

### Microbiome Processing

Microbiome analysis was completed on the distal colon samples. The DNA was extracted using the QIAamp PowerFecal Pro DNA kit (Qiagen; Cat No 51804) with an additional wash step to increase DNA purity. Extracted DNA was normalized using a Nanodrop 1000 spectrophotometer and the V4-V5 region of the 16S ribosomal DNA was amplified using 515FB and 926R primers attached to the Illumina adapter overhang. Samples were sequenced by the Integrated Microbiome Resource (Dalhousie University, Halifax, Nova Scotia, Canada) according to their protocol (Comeau, Douglas and Langille, 2017).

### Bioinformatic Analysis

Post-sequencing analyses were performed using the QIIME 2 platform (version 2021.4) (Bolyen *et al.*, 2019). Demultiplexed reads from two MiSeq runs were imported into the QIIME 2 environment and primers removed using the q2-cutadapt plugin (Martin, 2011). Quality control entailing of filtering, dereplication, chimera removal, denoising, and merging of paired-end reads were performed on each run separately using the DADA2 plugin with default settings (Callahan *et al.*, 2016). The resulting amplicon sequence variant (ASV) tables were merged for downstream analysis. A phylogenetic tree was constructed using a SATé-enabled phylogenetic placement (SEPP) technique as implemented in the q2-fragment-insertion plugin (Janssen *et al.*, 2018) using a backbone tree built from the Greengenes reference database (version 13.8) (McDonald *et al.*, 2012). For taxonomic classification, we trained a classifier on the full length 16S region and additionally incorporated environment-specific abundances weights specific to animal distal gut environment acquired from the *readytowear* repository (https://github.com/BenKaehler/readytowear). This weighted bespoke approach for taxonomic classification has been shown to significantly improve accuracy over common Naive Bayes classification methods (Kaehler *et al.*, 2019). Prior to diversity analysis, all ASVs that were not classified at least at the phylum level were discarded as contaminants and only samples with at least 1,000 sequences remaining were kept. A final table consisting of 7 CO, 8 MD, 9 MF, and 14 OO samples were retained for downstream analysis. Alpha-diversity metrics (Shannon’s diversity index, Faith’s phylogenetic diversity, ASV richness and Pielou’s Evenness) (Bolyen *et al.*, 2019). For beta diversity, we used the q2-DEICODE plug-in (Martino *et al.*, 2019) which calculates a form of Aitchison distances that is robust to high levels of sparsity, is compositionally aware, and circumvents the need for rarefying. Using the *q2-beta group significance* plugin, a permutational multivariate analysis of variance (PERMANOVA) test (α◻=◻0.05, with 999 permutations) was run on the robust Aitchison distances to determine differences between diet groups. The PERMANOVA test’s assumption of multivariate dispersion was assessed using a PERMDISP test. We utilized multinomial regression using Songbird (Morton *et al.*, 2019) to obtain ASV rankings most associated with each group. These differential ranks were visualized using Qurro (Fedarko *et al.*, 2020) and the differentials for the top 10 ASVs associated with each group exported into R (R Development Core Team, 2019) using the *qiime2R* package [https://github.com/jbisanz/qiime2R] for further custom visualization and statistical analysis.

### Statistical analysis

Statistical analyses were performed using R statistical software (R Development Core Team, 2019) and GraphPad Prism 9 (GraphPad Software, San Diego, California USA, www.graphpad.com) with *P* values below 0.05 considered statistically significant. The results are expressed as the mean value with standard error of the mean (SEM). When comparing diet groups, the Kruskal–Wallis test with the Dunn post hoc test was performed for nonparametric data, unless otherwise indicated. For bacterial differential abundance analysis, the log ratio of the top 5 ASVs associated with each diet group to top 5 ASV associated with MD were calculated and compared across groups using an ANOVA test. The false discovery rate resulting from multiple testing was controlled using the Benjamini-Hochberg method. The violin boxplots with jitters were produced using the ggplot2 (https://cran.r-project.org/web/packages/ggplot2/citation.html) package and further augmented with the ggpubr (https://rpkgs.datanovia.com/ggpubr/) package.

## References

Abulizi, N., Quin, C., Brown, K., Chan, Y.K., Gill, S.K. and Gibson, D.L. (2019) “Gut mucosal proteins and bacteriome are shaped by the saturation index of dietary lipids,” Nutrients, 11(2). doi:10.3390/nu11020418.

Al-Haddad, S. and Riddell, R.H. (2005) “The role of eosinophils in inflammatory bowel disease,” Gut, pp. 1674–1675. doi:10.1136/gut.2005.072595.

Ananthakrishnan, A.N., Khalili, H., Konijeti, G.G., Higuchi, L.M., de Silva, P., Fuchs, C.S., Willett, W.C., Richter, J.M. and Chan, A.T. (2014) “Long-term intake of dietary fat and risk of ulcerative colitis and Crohn’s disease,” Gut, 63(5), pp. 776–84. doi:10.1136/gutjnl-2013-305304.

Astrup, A., Magkos, F., Bier, D.M., Brenna, J.T., de Oliveira Otto, M.C., Hill, J.O., King, J.C., Mente, A., Ordovas, J.M., Volek, J.S., Yusuf, S. and Krauss, R.M. (2020) “Saturated Fats and Health: A Reassessment and Proposal for Food-Based Recommendations: JACC State-of-the-Art Review,” Journal of the American College of Cardiology, pp. 844–857. doi:10.1016/j.jacc.2020.05.077.

Bach-Faig, A., Berry, E.M., Lairon, D., Reguant, J., Trichopoulou, A., Dernini, S., Medina, F.X., Battino, M., Belahsen, R., Miranda, G. and Serra-Majem, L. (2011) “Mediterranean diet pyramid today. Science and cultural updates.,” Public health nutrition, 14(12A), pp. 2274–2284. doi:10.1017/S1368980011002515.

Barnett, J.A. and Gibson, D.L. (2020) “Separating the Empirical Wheat From the Pseudoscientific Chaff: A Critical Review of the Literature Surrounding Glyphosate, Dysbiosis and Wheat-Sensitivity,” Frontiers in Microbiology. doi:10.3389/fmicb.2020.556729.

Baxter, N.T., Zackular, J.P., Chen, G.Y. and Schloss, P.D. (2014) “Structure of the gut microbiome following colonization with human feces determines colonic tumor burden,” Microbiome, 2(1). doi:10.1186/2049-2618-2-20.

Bergstrom, K.S.B.B., Kissoon-Singh, V., Gibson, D.L., Ma, C., Montero, M., Sham, H.P., Ryz, N., Huang, T., Velcich, A., Finlay, B.B., Chadee, K. and Vallance, B.A. (2010) “Muc2 protects against lethal infectious colitis by disassociating pathogenic and commensal bacteria from the colonic mucosa,” PLoS Pathogens, 6(5), p. e1000902. doi:10.1371/journal.ppat.1000902.

Berry, D., Kuzyk, O., Rauch, I., Heider, S., Schwab, C., Hainzl, E., Decker, T., Müller, M., Strobl, B., Schleper, C., Urich, T., Wagner, M., Kenner, L. and Loy, A. (2015) “Intestinal microbiota signatures associated with inflammation history in mice experiencing recurring colitis,” Frontiers in Microbiology [Preprint]. doi:10.3389/fmicb.2015.01408.

Berry, D., Schwab, C., Milinovich, G., Reichert, J., ben Mahfoudh, K., Decker, T., Engel, M., Hai, B., Hainzl, E., Heider, S., Kenner, L., Müller, M., Rauch, I., Strobl, B., Wagner, M., Schleper, C., Urich, T. and Loy, A. (2012) “Phylotype-level 16S rRNA analysis reveals new bacterial indicators of health state in acute murine colitis,” ISME Journal, 6(11). doi:10.1038/ismej.2012.39.

Bloom, S.M., Bijanki, V.N., Nava, G.M., Sun, L., Malvin, N.P., Donermeyer, D.L., Dunne, W.M., Allen, P.M. and Stappenbeck, T.S. (2011) “Commensal Bacteroides species induce colitis in host-genotype-specific fashion in a mouse model of inflammatory bowel disease,” Cell Host and Microbe, 9(5), pp. 390–403. doi:10.1016/j.chom.2011.04.009.

Bloomfield, H.E., Koeller, E., Greer, N., MacDonald, R., Kane, R. and Wilt, T.J. (2016) “Effects on health outcomes of a mediterranean diet with no restriction on fat intake: A systematic review and meta-analysis,” Annals of Internal Medicine [Preprint]. doi:10.7326/M16-0361.

Bolyen, E., Rideout, J.R., Dillon, M.R., Bokulich, N.A., Abnet, C.C., Al-Ghalith, G.A., Alexander, H., Alm, E.J., Arumugam, M., Asnicar, F., Bai, Y., Bisanz, J.E., Bittinger, K., Brejnrod, A., Brislawn, C.J., Brown, C.T., Callahan, B.J., Caraballo-Rodríguez, A.M., Chase, J., Cope, E.K., da Silva, R., Diener, C., Dorrestein, P.C., Douglas, G.M., Durall, D.M., Duvallet, C., Edwardson, C.F., Ernst, M., Estaki, M., Fouquier, J., Gauglitz, J.M., Gibbons, S.M., Gibson, D.L., Gonzalez, A., Gorlick, K., Guo, J., Hillmann, B., Holmes, S., Holste, H., Huttenhower, C., Huttley, G.A., Janssen, S., Jarmusch, A.K., Jiang, L., Kaehler, B.D., Kang, K. bin, Keefe, C.R., Keim, P., Kelley, S.T., Knights, D., Koester, I., Kosciolek, T., Kreps, J., Langille, M.G.I., Lee, J., Ley, R., Liu, Y.X., Loftfield, E., Lozupone, C., Maher, M., Marotz, C., Martin, B.D., McDonald, D., McIver, L.J., Melnik, A. v., Metcalf, J.L., Morgan, S.C., Morton, J.T., Naimey, A.T., Navas-Molina, J.A., Nothias, L.F., Orchanian, et al. (2019) “Reproducible, interactive, scalable and extensible microbiome data science using QIIME 2,” Nature Biotechnology. Nature Publishing Group, pp. 852–857. doi:10.1038/s41587-019-0209-9.

Butera, A., di Paola, M., Pavarini, L., Strati, F., Pindo, M., Sanchez, M., Cavalieri, D., Boirivant, M. and de Filippo, C. (2018) “Nod2 Deficiency in mice is Associated with Microbiota Variation Favouring the Expansion of mucosal CD4+ LAP+ Regulatory Cells,” Scientific Reports, 8(1). doi:10.1038/s41598-018-32583-z.

Callahan, B.J., McMurdie, P.J., Rosen, M.J., Han, A.W., Johnson, A.J.A. and Holmes, S.P. (2016) “DADA2: High-resolution sample inference from Illumina amplicon data,” Nature Methods. doi:10.1038/nmeth.3869.

Christ, A., Lauterbach, M. and Latz, E. (2019) “Western Diet and the Immune System: An Inflammatory Connection,” Immunity, pp. 794–811. doi:10.1016/j.immuni.2019.09.020.

Comeau, A.M., Douglas, G.M. and Langille, M.G.I. (2017) “Microbiome Helper: a Custom and Streamlined Workflow for Microbiome Research,” mSystems doi:10.1128/msystems.00127-16.

Cooper, H.S., Murthy, S.N.S., Shah, R.S. and Sedergran, D.J. (1993) “Clinicopathologic study of dextran sulfate sodium experimental murine colitis,” Laboratory Investigation. doi:10.1016/S0021-5198(19)41298-5.

Couper, K.N., Blount, D.G. and Riley, E.M. (2008) “IL-10: The Master Regulator of Immunity to Infection,” The Journal of Immunology [Preprint]. doi:10.4049/jimmunol.180.9.5771.

Cranford, T.L., Enos, R.T., Velázquez, K.T., McClellan, J.L., Davis, J.M., Singh, U.P., Nagarkatti, M., Nagarkatti, P.S., Robinson, C.M. and Murphy, E.A. (2016) “Role of MCP-1 on inflammatory processes and metabolic dysfunction following high-fat feedings in the FVB/N strain,” International Journal of Obesity, 40(5), pp. 844–851. doi:10.1038/ijo.2015.244.

Crost, E.H., Tailford, L.E., Le Gall, G., Fons, M., Henrissat, B. and Juge, N. (2013) “Utilisation of Mucin Glycans by the Human Gut Symbiont Ruminococcus gnavus Is Strain-Dependent,” PLoS ONE, 8(10). doi:10.1371/journal.pone.0076341.

DeCoffe, D., Quin, C., Gill, S.K., Tasnim, N., Brown, K., Godovannyi, A., Dai, C., Abulizi, N., Chan, Y.K., Ghosh, S. and Gibson, D.L. (2016) “Dietary lipid type, rather than total number of calories, alters outcomes of enteric infection in mice,” Journal of Infectious Diseases, 213(11), pp. 1846–1856. doi:10.1093/infdis/jiw084.

Fedarko, M.W., Martino, C., Morton, J.T., González, A., Rahman, G., Marotz, C.A., Minich, J.J., Allen, E.E. and Knight, R. (2020) “Visualizing ‘omic feature rankings and log-ratios using Qurro,” NAR Genomics and Bioinformatics, 2(2). doi:10.1093/nargab/lqaa023.

Feng, Q., Liang, S., Jia, H., Stadlmayr, A., Tang, L., Lan, Z., Zhang, D., Xia, H., Xu, Xiaoying, Jie, Z., Su, L., Li, Xiaoping, Li, Xin, Li, J., Xiao, L., Huber-Schönauer, U., Niederseer, D., Xu, Xun, Al-Aama, J.Y., Yang, H., Wang, Jian, Kristiansen, K., Arumugam, M., Tilg, H., Datz, C. and Wang, Jun (2015) “Gut microbiome development along the colorectal adenoma-carcinoma sequence,” Nature Communications, 6. doi:10.1038/ncomms7528.

Forbes, A., Escher, J., Hébuterne, X., Kłęk, S., Krznaric, Z., Schneider, S., Shamir, R., Stardelova, K., Wierdsma, N., Wiskin, A.E. and Bischoff, S.C. (2017) “ESPEN guideline: Clinical nutrition in inflammatory bowel disease,” Clinical Nutrition. doi:10.1016/j.clnu.2016.12.027.

Gálvez, J. (2014) “Role of Th17 Cells in the Pathogenesis of Human IBD,” ISRN Inflammation, 2014, pp. 1–14. doi:10.1155/2014/928461.

Ganesh, B.P., Klopfleisch, R., Loh, G. and Blaut, M. (2013) “Commensal Akkermansia muciniphila Exacerbates Gut Inflammation in Salmonella Typhimurium-Infected Gnotobiotic Mice,” PLoS ONE, 8(9). doi:10.1371/journal.pone.0074963.

Haskey, N. and Gibson, D.L. (2017) “An examination of diet for the maintenance of remission in inflammatory bowel disease,” Nutrients, 9(3). doi:10.3390/nu9030259.

Hibberd, A.A., Lyra, A., Ouwehand, A.C., Rolny, P., Lindegren, H., Cedgård, L. and Wettergren, Y. (2017) “Intestinal microbiota is altered in patients with colon cancer and modified by probiotic intervention,” BMJ Open Gastroenterology, 4(1). doi:10.1136/bmjgast-2017-000145.

Hou, J.K., Abraham, B. and El-Serag, H. (2011) “Dietary intake and risk of developing inflammatory bowel disease: a systematic review of the literature.,” American Journal of Gastroenterology, 106(4), pp. 563–573. doi:10.1038/ajg.2011.44.

Ikemoto, S., Takahashi, M., Tsunoda, N., Maruyama, K., Itakura, H. and Ezaki, O. (1996) “High-fat diet-induced hyperglycemia and obesity in mice: Differential effects of dietary oils,” Metabolism: Clinical and Experimental, 45(12), pp. 1539–1546. doi:10.1016/S0026-0495(96)90185-7.

Imam, T., Park, S., Kaplan, M.H. and Olson, M.R. (2018) “Effector T helper cell subsets in inflammatory bowel diseases,” Frontiers in Immunology. doi:10.3389/fimmu.2018.01212.

Janssen, S., McDonald, D., Gonzalez, A., Navas-Molina, J.A., Jiang, L., Xu, Z.Z., Winker, K., Kado, D.M., Orwoll, E., Manary, M., Mirarab, S. and Knight, R. (2018) “Phylogenetic Placement of Exact Amplicon Sequences Improves Associations with Clinical Information,” mSystems. doi:10.1128/msystems.00021-18.

Kaehler, B.D., Bokulich, N.A., McDonald, D., Knight, R., Caporaso, J.G. and Huttley, G.A. (2019) “Species abundance information improves sequence taxonomy classification accuracy,” Nature Communications, 10(1). doi:10.1038/s41467-019-12669-6.

Kim, J.J., Shajib, M.S., Manocha, M.M. and Khan, W.I. (2012) “Investigating intestinal inflammation in DSS-induced model of IBD,” Journal of Visualized Experiments, 60, pp. 1–6. doi:10.3791/3678.

Laffin, M., Fedorak, R., Zalasky, A., Park, H., Gill, A., Agrawal, A., Keshteli, A., Hotte, N. and Madsen, K.L. (2019) “A high-sugar diet rapidly enhances susceptibility to colitis via depletion of luminal short-chain fatty acids in mice,” Scientific Reports, 9(1). doi:10.1038/s41598-019-48749-2.

Lallès, J.P. (2010) “Intestinal alkaline phosphatase: Multiple biological roles in maintenance of intestinal homeostasis and modulation by diet,” Nutrition Reviews. doi:10.1111/j.1753-4887.2010.00292.x.

Lee, J.C., Lee, H.Y., Kim, T.K., Kim, M.S., Park, Y.M., Kim, J., Park, K., Kweon, M.N., Kim, S.H., Bae, J.W., Hur, K.Y. and Lee, M.S. (2017) “Obesogenic diet-induced gut barrier dysfunction and pathobiont expansion aggravate experimental colitis,” PLoS ONE, 12(11). doi:10.1371/journal.pone.0187515.

Li, T., Qiu, Y., Yang, H.S., Li, M.Y., Zhuang, X.J., Zhang, S.H., Feng, R., Chen, B.L., He, Y., Zeng, Z.R. and Chen, M.H. (2020) “Systematic review and meta-analysis: Association of a pre-illness Western dietary pattern with the risk of developing inflammatory bowel disease,” Journal of Digestive Diseases, 21(7), pp. 362–371. doi:10.1111/1751-2980.12910.

Li, Xue, Wei, X., Sun, Y., Du, J., Li, Xin, Xun, Z. and Li, Y.C. (2019) “High-fat diet promotes experimental colitis by inducing oxidative stress in the colon,” American Journal of Physiology - Gastrointestinal and Liver Physiology, 317(4). doi:10.1152/ajpgi.00103.2019.

Li, Y., Dong, J., Xiao, H., Zhang, S., Wang, B., Cui, M. and Fan, S. (2020) “Gut commensal derived-valeric acid protects against radiation injuries,” Gut Microbes, 11(4), pp. 789–806. doi:10.1080/19490976.2019.1709387.

Lindemans, C.A., Calafiore, M., Mertelsmann, A.M., O’Connor, M.H., Dudakov, J.A., Jenq, R.R., Velardi, E., Young, L.F., Smith, O.M., Lawrence, G., Ivanov, J.A., Fu, Y.Y., Takashima, S., Hua, G., Martin, M.L., O’Rourke, K.P., Lo, Y.H., Mokry, M., Romera-Hernandez, M., Cupedo, T., Dow, L.E., Nieuwenhuis, E.E., Shroyer, N.F., Liu, C., Kolesnick, R., van den Brink, M.R.M. and Hanash, A.M. (2015) “Interleukin-22 promotes intestinal-stem-cell-mediated epithelial regeneration,” Nature, 528(7583), pp. 560–564. doi:10.1038/nature16460.

Livanos, A.E., Greiner, T.U., Vangay, P., Pathmasiri, W., Stewart, D., McRitchie, S., Li, H., Chung, J., Sohn, J., Kim, S., Gao, Z., Barber, C., Kim, J., Ng, S., Rogers, A.B., Sumner, S., Zhang, X.S., Cadwell, K., Knights, D., Alekseyenko, A., B’ckhed, F. and Blaser, M.J. (2016) “Antibiotic-mediated gut microbiome perturbation accelerates development of type 1 diabetes in mice,” Nature Microbiology, 1. doi:10.1038/nmicrobiol.2016.140.

Loonen, L.M.P., Stolte, E.H., Jaklofsky, M.T.J., Meijerink, M., Dekker, J., van Baarlen, P. and Wells, J.M. (2014) “REG3γ-deficient mice have altered mucus distribution and increased mucosal inflammatory responses to the microbiota and enteric pathogens in the ileum,” Mucosal Immunology, 7(4). doi:10.1038/mi.2013.109.

Madsen, K.L., Malfair, D., Gray, D., Doyle, J.S., Jewell, L.D. and Fedorak, R.N. (1999) “Interleukin-10 gene-deficient mice develop a primary intestinal permeability defect in response to enteric microflora,” Inflammatory Bowel Diseases, 5(4), pp. 262–270. doi:10.1097/00054725-199911000-00004.

Martin, M. (2011) “Cutadapt removes adapter sequences from high-throughput sequencing reads,” EMBnet.journal, 17(1). doi:10.14806/ej.17.1.200.

Martino, C., Morton, J.T., Marotz, C.A., Thompson, L.R., Tripathi, A., Knight, R. and Zengler, K. (2019) “A Novel Sparse Compositional Technique Reveals Microbial Perturbations,” mSystems, 4(1). doi:10.1128/msystems.00016-19.

McDonald, D., Price, M.N., Goodrich, J., Nawrocki, E.P., Desantis, T.Z., Probst, A., Andersen, G.L., Knight, R. and Hugenholtz, P. (2012) “An improved Greengenes taxonomy with explicit ranks for ecological and evolutionary analyses of bacteria and archaea,” ISME Journal, 6(3). doi:10.1038/ismej.2011.139.

Meshkibaf, S., Martins, A.J., Henry, G.T. and Kim, S.O. (2016) “Protective role of G-CSF in dextran sulfate sodium-induced acute colitis through generating gut-homing macrophages,” Cytokine, 78, pp. 69–78. doi:10.1016/j.cyto.2015.11.025.

Morampudi, V., Dalwadi, U., Bhinder, G., Sham, H.P., Gill, S.K., Chan, J., Bergstrom, K.S.B., Huang, T., Ma, C., Jacobson, K., Gibson, D.L. and Vallance, B.A. (2016) “The goblet cell-derived mediator RELM-β drives spontaneous colitis in Muc2-deficient mice by promoting commensal microbial dysbiosis,” Mucosal Immunology, 9(5), pp. 1218–1233. doi:10.1038/mi.2015.140.

Morton, J.T., Marotz, C., Washburne, A., Silverman, J., Zaramela, L.S., Edlund, A., Zengler, K. and Knight, R. (2019) “Establishing microbial composition measurement standards with reference frames,” Nature Communications, 10(1). doi:10.1038/s41467-019-10656-5.

Neurath, M.F. (2014) “Cytokines in inflammatory bowel disease,” Nature Reviews Immunology doi:10.1038/nri3661.

Ng, S.C., Shi, H.Y., Hamidi, N., Underwood, F.E., Tang, W., Benchimol, E.I., Panaccione, R., Ghosh, S., Wu, J.C.Y., Chan, F.K.L., Sung, J.J.Y. and Kaplan, G.G. (2017) “Worldwide incidence and prevalence of inflammatory bowel disease in the 21st century: a systematic review of population-based studies,” The Lancet, 390(10114), pp. 2769–2778. doi:10.1016/S0140-6736(17)32448-0.

Nunes, T., Bernardazzi, C. and de Souza, H.S. (2014) “Cell death and inflammatory bowel diseases: Apoptosis, necrosis, and autophagy in the intestinal epithelium,” BioMed Research International. doi:10.1155/2014/218493.

Opferman, J.T. and Korsmeyer, S.J. (2003) “Apoptosis in the development and maintenance of the immune system,” Nature Immunology. doi:10.1038/ni0503-410.

Parada Venegas, D., de la Fuente, M.K., Landskron, G., González, M.J., Quera, R., Dijkstra, G., Harmsen, H.J.M., Faber, K.N. and Hermoso, M.A. (2019) “Short Chain Fatty Acids (SCFAs)-Mediated Gut Epithelial and Immune Regulation and Its Relevance for Inflammatory Bowel Diseases.,” Frontiers in immunology, 10(277). doi:10.3389/fimmu.2019.00277.

Patterson, E., O’ Doherty, R.M., Murphy, E.F., Wall, R., O’ Sullivan, O., Nilaweera, K., Fitzgerald, G.F., Cotter, P.D., Ross, R.P. and Stanton, C. (2014) “Impact of dietary fatty acids on metabolic activity and host intestinal microbiota composition in C57BL/6J mice.,” The British journal of nutrition, 111(11), pp. 1–13. doi:10.1017/S0007114514000117.

Paz-Filho, G., Wong, M.-L., Licinio, J. and Mastronardi, C. (2012) “Leptin therapy, insulin sensitivity, and glucose homeostasis,” Indian Journal of Endocrinology and Metabolism, 16(9). doi:10.4103/2230-8210.105571.

R Development Core Team, R. (2019) R: A Language and Environment for Statistical Computing, R Foundation for Statistical Computing. doi:10.1007/978-3-540-74686-7.

Rabiee, A., Galiatsatos, P., Salas-Carrillo, R., Thompson, M.J., Andersen, D.K. and Elahi, D. (2011) “Pancreatic polypeptide administration enhances insulin sensitivity and reduces the insnlin requirement of patients on insulin pump therapy,” Journal of Diabetes Science and Technology, 5(6). doi:10.1177/193229681100500629.

Ramasamy, S., Nguyen, D.D., Eston, M.A., Nasrin Alam, S., Moss, A.K., Ebrahimi, F., Biswas, B., Mostafa, G., Chen, K.T., Kaliannan, K., Yammine, H., Narisawa, S., Millán, J.L., Warren, H.S., Hohmann, E.L., Mizoguchi, E., Reinecker, H.C., Bhan, A.K., Snapper, S.B., Malo, M.S. and Hodin, R.A. (2011) “Intestinal alkaline phosphatase has beneficial effects in mouse models of chronic colitis,” Inflammatory Bowel Diseases, 17(2). doi:10.1002/ibd.21377.

Ramos, G.P. and Papadakis, K.A. (2019) “Mechanisms of Disease: Inflammatory Bowel Diseases,” Mayo Clinic Proceedings [Preprint]. doi:10.1016/j.mayocp.2018.09.013.

Rath, H.C., Herfarth, H.H., Ikeda, J.S., Grenther, W.B., Hamm, T.E., Balish, E., Taurog, J.D., Hammer, R.E., Wilson, K.H. and Sartor, R.B. (1996) “Normal luminal bacteria, especially bacteroides species, mediate chronic colitis, gastritis, and arthritis in HLA-B27/human β2 microglobulin transgenic rats,” Journal of Clinical Investigation [Preprint]. doi:10.1172/JCI118878.

Sasson, A.N., Ingram, R.J.M., Zhang, Z., Taylor, L.M., Ananthakrishnan, A.N., Kaplan, G.G., Ng, S.C., Ghosh, S. and Raman, M. (2021) “The role of precision nutrition in the modulation of microbial composition and function in people with inflammatory bowel disease,” The Lancet Gastroenterology & Hepatology, S2468–1253. doi:10.1016/S2468-1253(21)00097-2.

Schwarz, M., Lund, E.G., Setchell, K.D.R., Kayden, H.J., Zerwekh, J.E., Björkhem, I., Herz, J. and Russell, D.W. (1996) “Disruption of cholesterol 7α-hydroxylase gene in mice. II. Bile acid deficiency is overcome by induction of oxysterol 7α-hydroxylase,” Journal of Biological Chemistry. doi:10.1074/jbc.271.30.18024.

Serra-Majem, L., Bes-Rastrollo, M., Román-Viñas, B., Pfrimer, K., Sánchez-Villegas, A. and Martínez-González, M.A. (2009) “Dietary patterns and nutritional adequacy in a Mediterranean country,” British Journal of Nutrition [Preprint]. doi:10.1017/S0007114509990559.

Sibai, M., Altuntaş, E., Ylldlrlm, B., Öztürk, G., Ylldlrlm, S. and Demircan, T. (2020) “Microbiome and Longevity: High Abundance of Longevity-Linked Muribaculaceae in the Gut of the Long-Living Rodent Spalax leucodon,” OMICS A Journal of Integrative Biology, 24(10). doi:10.1089/omi.2020.0116.

Sonnenburg, J.L. and Sonnenburg, E.D. (2019) “Vulnerability of the industrialized microbiota,” Science. doi:10.1126/science.aaw9255.

Stagg, A.J. (2018) “Intestinal Dendritic Cells in Health and Gut Inflammation,” Frontiers in immunology, p. 2883. doi:10.3389/fimmu.2018.02883.

Stidham, R.W. and Higgins, P.D.R. (2018) “Colorectal Cancer in Inflammatory Bowel Disease,” Clinics in Colon and Rectal Surgery, 31(3), pp. 168–178. doi:10.1055/s-0037-1602237.

Teixeira, L.G., Leonel, A.J., Aguilar, E.C., Batista, N. v., Alves, A.C., Coimbra, C.C., Ferreira, A.V.M., de Faria, A.M.C., Cara, D.C. and Alvarez Leite, J.I. (2011) “The combination of high-fat diet-induced obesity and chronic ulcerative colitis reciprocally exacerbates adipose tissue and colon inflammation,” Lipids in Health and Disease, 10. doi:10.1186/1476-511X-10-204.

Tjonneland, A., Overvad, K., Bergmann, M.M., Nagel, G., Linseisen, J., Hallmans, G., Palmqvist, R., Sjodin, H., Hagglund, G., Berglund, G., Lindgren, S., Grip, O., Palli, D., Day, N.E., Khaw, K.-T., Bingham, S., Riboli, E., Kennedy, H. and Hart, A. (2009) “Linoleic acid, a dietary n-6 polyunsaturated fatty acid, and the aetiology of ulcerative colitis: a nested case-control study within a European prospective cohort study.,” Gut, 58(12), pp. 1606–11. doi:10.1136/gut.2008.169078.

Verdugo-Meza, A., Ye, J., Dadlani, H., Ghosh, S. and Gibson, D.L. (2020) “Connecting the dots between inflammatory bowel disease and metabolic syndrome: A focus on gut-derived metabolites,” Nutrients. doi:10.3390/nu12051434.

Visconti, A., le Roy, C.I., Rosa, F., Rossi, N., Martin, T.C., Mohney, R.P., Li, W., de Rinaldis, E., Bell, J.T., Venter, J.C., Nelson, K.E., Spector, T.D. and Falchi, M. (2019) “Interplay between the human gut microbiome and host metabolism,” Nature Communications, 10(1). doi:10.1038/s41467-019-12476-z.

de Vos, W.M. (2017) “Microbe profile: Akkermansia muciniphila: A conserved intestinal symbiont that acts as the gatekeeper of our mucosa,” Microbiology (United Kingdom), 163(5). doi:10.1099/mic.0.000444.

Weigmann, B., Tubbe, I., Seidel, D., Nicolaev, A., Becker, C. and Neurath, M.F. (2007) “Isolation and subsequent analysis of murine lamina propria mononuclear cells from colonic tissue,” Nature Protocols, 2(10), pp. 2307–2311. doi:10.1038/nprot.2007.315.

Williams, I.R. and Parkos, C.A. (2007) “Colonic Neutrophils in Inflammatory Bowel Disease: Double-Edged Swords of the Innate Immune System With Protective and Destructive Capacity,” Gastroenterology. doi:10.1053/j.gastro.2007.10.031.

Wong, C.K., Botta, A., Pither, J., Dai, C., Gibson, W.T. and Ghosh, S. (2015) “A high-fat diet rich in corn oil reduces spontaneous locomotor activity and induces insulin resistance in mice,” Journal of Nutritional Biochemistry [Preprint]. doi:10.1016/j.jnutbio.2014.11.004.

Ye, J., Haskey, N., Dadlani, H., Zubaidi, H., Barnett, J.A., Ghosh, S. and Gibson, D.L. (2021) “Deletion of mucin 2 induces colitis with concomitant metabolic abnormalities in mice,” American Journal of Physiology - Gastrointestinal and Liver Physiology, 320(5), pp. G791–G803. doi:10.1152/AJPGI.00277.2020.

