## Supplemental Materials for "A Mediterranean-like fat blend protects against the development of severe colitis in the mucin-2 deficient murine model" (from another version of this preprint: version 2, 2021-10-04)

**Supplemental Information**

**Table S1. Nutrient composition of experimental diets.** Four diets were examined in this study – Mediterranean Diet, Olive Oil, Corn Oil and Anhydrous Milk Fat. All diets were isocaloric, iso-nitrogenous with similar protein, carbohydrate, vitamin and mineral profiles. Lipid content was altered to reflect the various fatty acid profiles.

|  | Mediterranean Diet  (TD. 170674) | Olive Oil (TD. 130128) | Corn Oil (TD. 120022) | Anhydrous Milk Fat  (TD. 120021) |
| --- | --- | --- | --- | --- |
| Energy Density (Kcal/g) | 4.5 | 4.5 | 4.5 | 4.5 |
| Protein (%) | 19 | 19 | 19 | 19 |
| Carbohydrates (%) | 40.1 | 40.1 | 40.1 | 40.1 |
| Fat (%) | 40.8 | 40.8 | 40.8 | 40.8 |
| Total SFA (g/kg) | 38.3 | 34.0 | 28.9 | 125.2 |
| Total MUFA (g/kg) | 136.6 | 139.7 | 54.6 | 61.8 |
| Total PUFA (g/kg) | 24.7 | 26.3 | 116.3 | 12.8 |
| n-6 PUFA:n3 PUFA ratio (n:1) | 2 | 2 | 46 | 5 |

***Figure S1. Related to Figure 1.* Type of fat does not impact body weight gain or food intake.** MD (n=32), Olive oil (n=33), Corn oil (n=30) and Milk fat diets (n=30) food intake (A) body weights (B) and clinical signs (C) were recorded weekly from weaning until the end of the study. Clinical scores shown in (C) are the final clinical scores at 12 weeks of age for Muc2**^-/-^** mice. Data represent means ± SEM. Two-way ANOVA was performed, a value of P <0.05 was considered statistically significant.

**B**

**A**

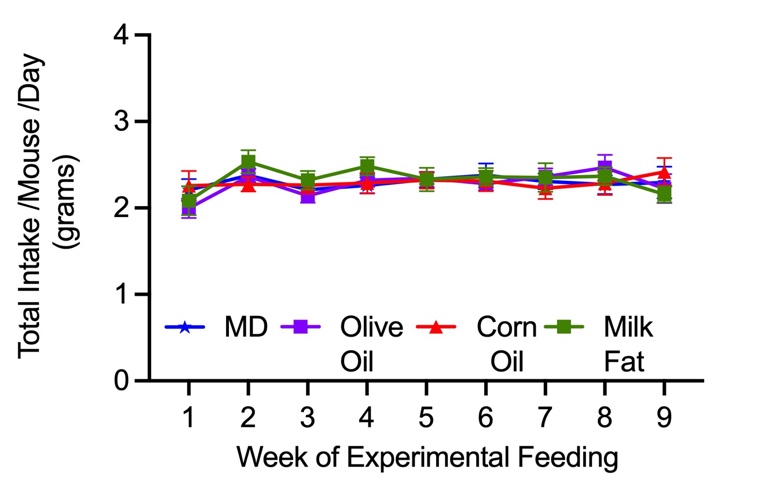

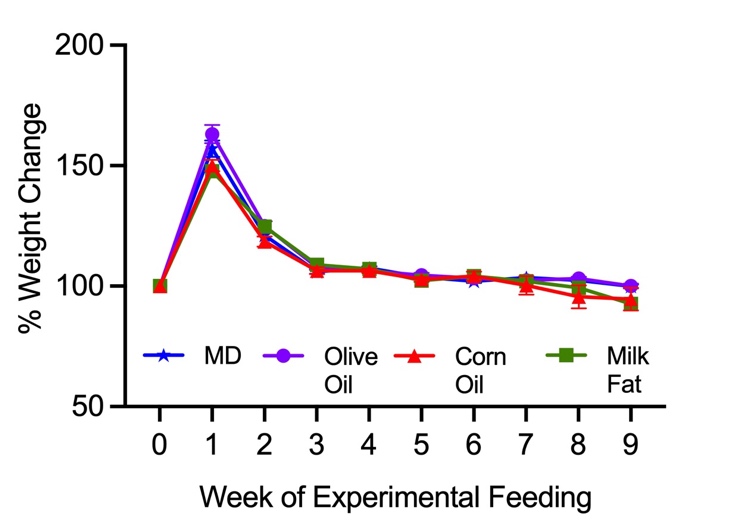

**C**

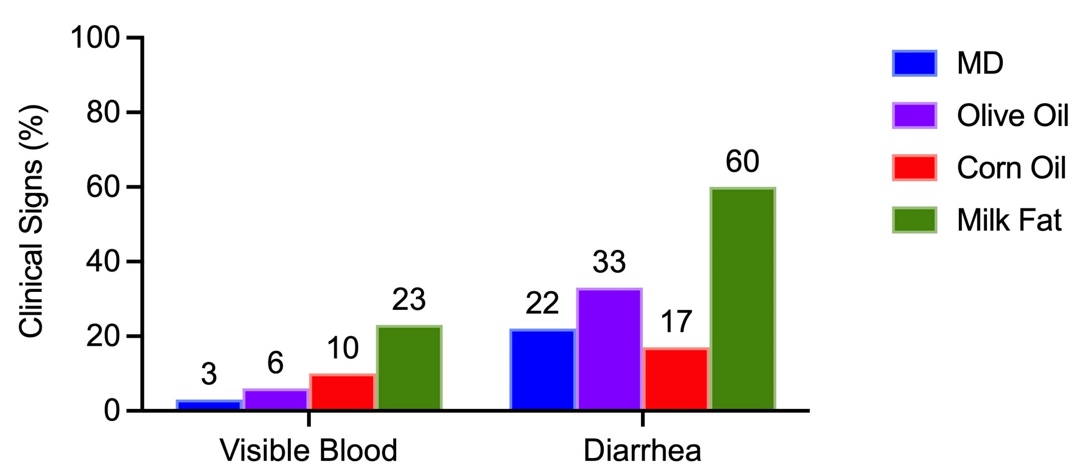

***Figure S2. Related to Figure 2.*** Type of fat impacts neutrophil infiltration in the Muc2^-/-^ mice fed Mediterranean Diet (n=10), Olive oil (n=16), Corn oil (n=12) and Milk fat diets (n= 16). Colonic tissue sections were incubated with a primary antibody rabbit polyclonal antibody-1 for myeloperoxidase and secondary antibody goat anti-rabbit IgG 488-conjugated antibody to visualize neutrophils. Data represent means ± SEM. A Kruskal-Wallis test was performed with Dunn's multiple comparisons test, a value of P <0.05 was considered statistically significant.

Mediterranean Diet

Olive Oil

**MPO+**

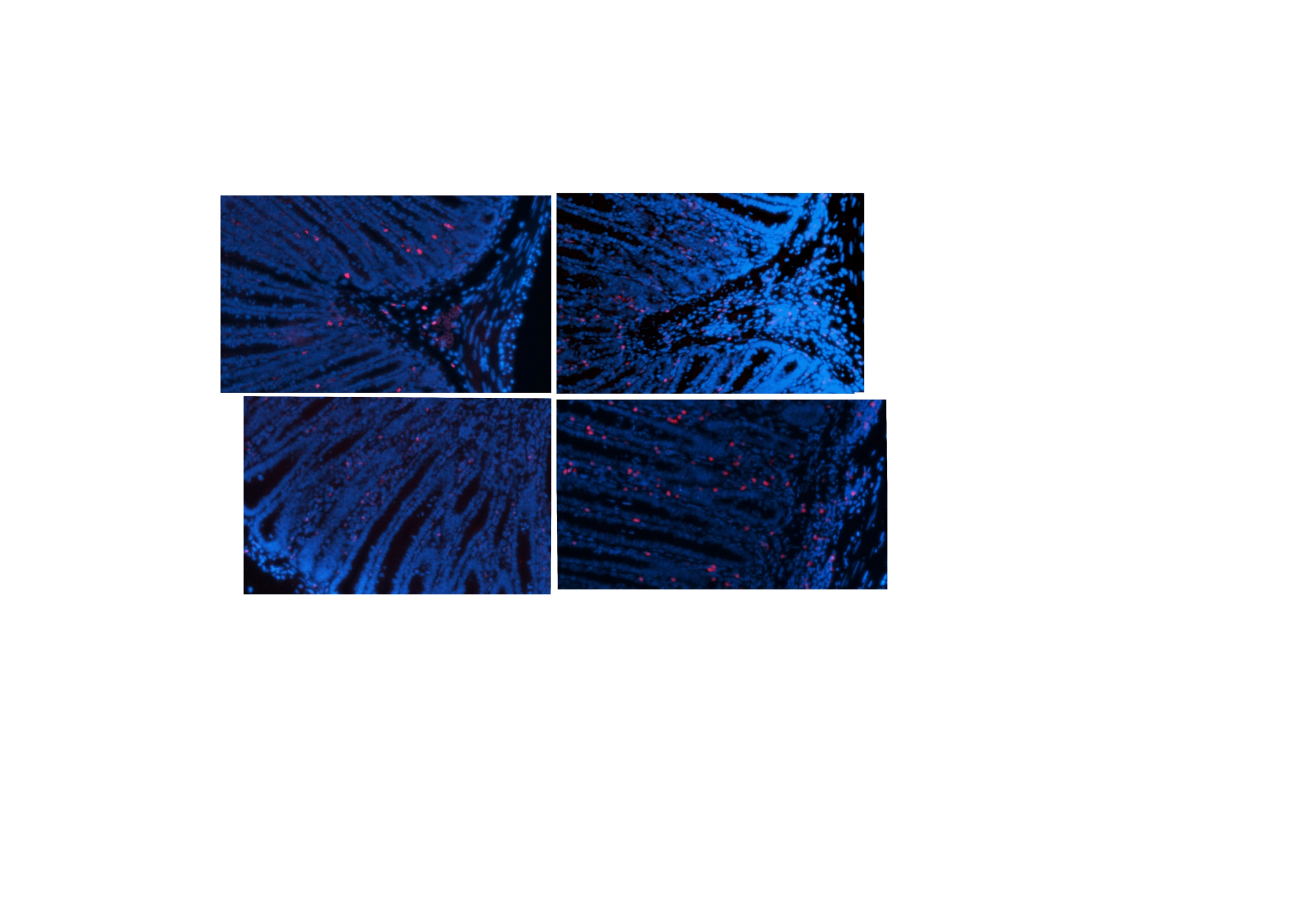

**100 um**

Corn Oil

Milk Fat

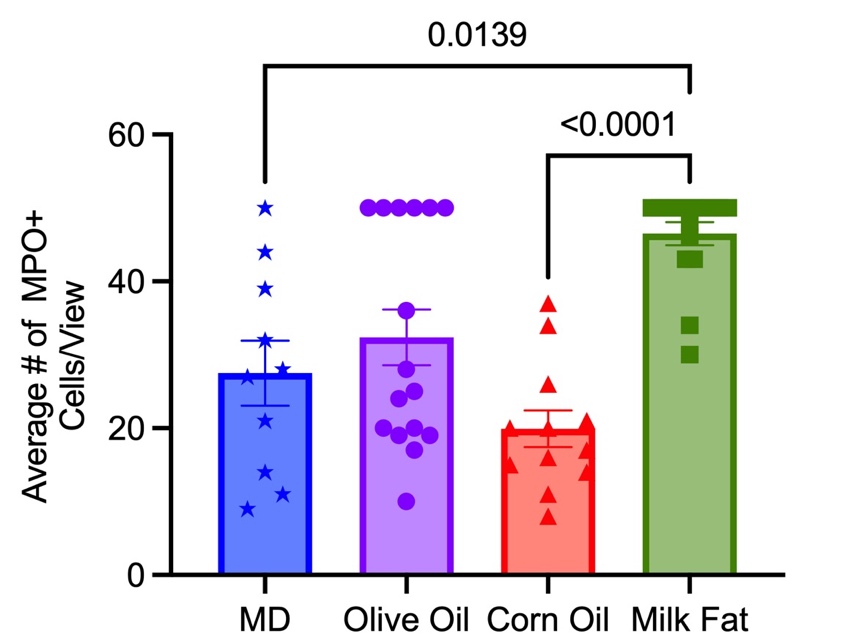

***Table S3. Related to Figure 3.*** Levels of Colonic mRNA Relative Gene Expression Encoding Inflammatory Mediators Among Muc2^-/-^ Mice Fed the MD (n=23), Olive Oil (n=14), Corn Oil (n=14), Milk Fat (n= 14).

| Relative Gene Expression,  Mean ± SEM, by Diet | | |
| --- | --- | --- |
| Response |  | *P* Value^a^ |
| TNF-α |  | 0.5546 |
| MD | 0.0389 ± 0.0093 |  |
| Olive Oil | 0.0213 ± 0.0042 |  |
| Corn Oil | 0.0242 ± 0.0098 |  |
| Milk Fat | 0.0644 ± 0.0241 |  |
| FOXP3 |  | 0.7702 |
| MD | 0.0536 ± 0.0094 |  |
| Olive Oil | 0.0576 ± 0.0125 |  |
| Corn Oil | 0.0577 ± 0.0083 |  |
| Milk Fat | 0.0677 ± 0.0144 |  |
| TGF-β1 |  | 0.5793 |
| MD | 0.0224 ± 0.0055 |  |
| Olive Oil | 0.0281 ± 0.0079 |  |
| Corn Oil | 0.0573 ± 0.0187 |  |
| Milk Fat | 0.0157 ± 0.0050 |  |
| Ebi3 |  | 0.6378 |
| MD | 2.51 ± 0.399 |  |
| Olive Oil | 2.38 ± 0.444 |  |
| Corn Oil | 2.96 ± 0.421 |  |
| Milk Fat | 2.23 ± 0.519 |  |
| IL-22 |  | 0.6169 |
| MD | 0.0218 ± 0.0039 |  |
| Olive Oil | 0.0296 ± 0.0061 |  |
| Corn Oil | 0.0392 ± 0.0151 |  |
| Milk Fat | 0.0877 ± 0.0331 |  |

Abbreviations: TNF-α: tumor necrosis factor alpha; FOXP3: forkhead box P3; TGF-β1:	transforming growth factor beta 1; Ebi3: Epstein-Barr Virus induced 3; IL-22: interleukin 22

^a^By the Kruskal-Wallis test. A value of P <0.05 was considered statistically significant.

**Table S3. *Related to Figure 3.*** Primer sequences used for qPCR analysis

| Primer | Forward | Reverse |
| --- | --- | --- |
| TBP | *accgtgaatcttggctgtaaac* | *accgtgaatcttggctgtaaac* |
| Reg3-γ | *cccgtataaccatcaccatcat* | *ggcatctttcttggcaacttc* |
| RELM-β | *atgggtgtcactggatgtgctt* | *agcactggcagtggcaagta* |
| TNF-α | *catcttctcaaaattcgagtgacaa* | *tgggagtagaacaaggtacaaccc* |
| TGF-β1 | *gaccgcaacaacgccatcta* | *agccctgtattccgtctcctt* |
| IL-1β | *agctatccttgtgcaagtgtc* | *cccttcatcttttggggtcc* |
| IL-6 | *gaggataccactcccaacagacc* | *aagtgcatcatcgttgttcataca* |
| IL-10 | *agggccctttgctatggtgt* | *tggccacagttttcagggat* |
| IL-17 | *tccgaggagtcagtgctaaa* | *tccgaggagtcagtgctaaa* |
| IL-22 | *agctcctgtcacatcagcg* | *agcttcttctcgctcagacg* |
| FOXP3 | *ggcccttctccaggacaga* | *ggcccttctccaggacaga* |
| GLP-1 | *cagaagttggtcgtgaggca* | *gcctttcaccagccaagcaa* |
| Ebi3 | *cccggacatcttctctct* | *gaggctccagtcacttg* |

Abbreviations: TBP: TATA-box binding protein; Reg3-γ: regenerating islet-derived protein 3 gamma; RELM-β: resistin-like molecule beta; TNF-α: tumor necrosis factor alpha; TGF-β1: transforming growth factor beta 1; IL: interleukin; FOXP3: forkhead box P3; GLP-1: glucagon-like peptide 1; Ebi3: Epstein-Barr Virus induced 3.

***Table S4. Related to Figure 4.*** Flow Cytometry Analysis of Muc2^-/-^ Mice Fed the MD (n=7), Olive Oil (n=13), Corn Oil (n=9) and Milk Fat Diets (n= 10).

| Flow Cytometry Analysis  Mean ± SEM, by Diet | | | | | |
| --- | --- | --- | --- | --- | --- |
|  | MD  (n=7) | Olive Oil (n=13) | Corn Oil  (n=9) | Milk Fat (n=10) | *P* Value^a^ |
| % CD45+ | 61.4 ±  5.82 | 69.8 ±  3.06 | 69.7 ±  3.66 | 74.9 ±  2.09 | 0.2033 |
| % Inflammatory Monocytes | 2.91 ±  0.614 | 4.70 ±  0.828 | 2.97 ±  0.694 | 4.64 ±  0.652 | 0.2224 |
| % Neutrophils | 0.680 ±  0.179 | 1.36 ±  0.205 | 0.974 ±  0.188 | 1.28 ±  0.276 | 0.2577 |
| % IFNγ Producing Cells | 11.3 ±  1.47 | 12.7 ±  1.38 | 8.45 ±  1.67 | 17.0 ±  1.77 | 0.0203 |
| % TNFα Producing  Cells | 4.76 ±  0.982 | 11.3 ±  1.70 | 4.10 ±  1.10 | 9.85 ±  1.44 | 0.0044  (Corn Oil vs Olive Oil  P =0.0153) |
| % Dendritic Cells  103+ | 55.4 ±  5.33 | 45.3 ±  2.35 | 50.1 ±  2.75^b,d^ | 44.1 ±  4.21^b,d^ | 0.2355 |
| % Dendritic Cells  CD11b+ | 15.2 ±  1.96 | 19.2 ±  1.72 | 15.7 ±  0.898^c^ | 14.8 ±  1.09^c^ | 0.6197 |
| % T helper  Cells | 16.6 ±  2.05 | 21.8 ±  1.64 | 17.5 ±  2.27 | 23.9 ±  1.69 | 0.0431  (MD vs Milk Fat  P =0.0063) |
| % Cytotoxic T Cells | 9.70 ±  1.17 | 7.16 ±  0.923^c^ | 8.35 ±  1.30^c^ | 6.11 ±  1.14^c^ | 0.1666 |
| % B cells | 25.9 ±  3.27 | 18.1 ±  1.40 | 18.7 ±  2.79 | 18.9 ±  1.47 | 0.1880 |

^a^ By the Kruskal-Wallis test with Dunn's multiple comparisons test. A value of P <0.05 was considered statistically significant.

^b^ One sample detected as an outlier.

^c^Three samples detected as outliers.

^d^ Non-detect in sample.

***Table S5. Related to Figure 5.*** Serum Cytokine Levels of Muc2^-/-^ Mice Fed the MD (n=7), Olive Oil (n=8), Corn oil (n=7) and Milk Fat Diets (n= 7).

| Serum Cytokines  Mean ± SEM, by Diet | | | | | |
| --- | --- | --- | --- | --- | --- |
|  | MD (n=7) | Olive Oil (n=8) | Corn Oil (n=7) | Milk Fat (n=7) | *P* Value^a^ |
| Eotaxin | 380 ± 45.6 | 315 ± 45.3 | 321 ±39.7 | 347 ± 37.3 | 0.6693 |
| GM-CSF | 16.5 ± 7.77 | 32.6 ±15.1 | 19.3 ± 7.35 | 16.2 ± 6.05 | 0.5163 |
| IL-1α | 273 ± 62.9 | 190 ± 47.0 | 229 ± 88.6 | 195 ± 53.6 | 0.7434 |
| IL-1β | 1.49 ± 0.251 | 1.36 ± 0.287 | 2.05 ± 0.669 | 3.06 ± 1.21 | 0.8667 |
| IL-2 | 10.6 ± 4.43 | 26.5 ± 16.8 | 13.0 ± 3.29 | 10.3 ± 3.53^b^ | 0.8158 |
| IL-3 | 2.09 ± 0.955^c^ | 2.28 ± 0.669 | 2.42 ± 0.728^b^ | 1.90 ± 0.842^d^ | 0.8610 |
| IL-4 | 1.06 ± 0.299 | 0.604 ± 0.113 | 0.997 ± 0.386 | 0.827 ± 0.363 | 0.7235 |
| 1L-7 | 12.3 ± 5.39 | 4.36 ± 1.09 | 4.64 ± 1.79 | 3.14 ± 1.38 | 0.3422 |
| 1L-9 | 83.1 ± 9.67 | 139 ± 31.6 | 82.7 ± 23.1 | 90.6 ± 29.6 | 0.5883 |
| 1L-12(P40) | 10.1 ± 3.82^b^ | 12.1 ± 3.16 | 7.28 ± 2.42^b^ | 6.94 ± 3.21^c^ | 0.4717 |
| 1L-12 (P70) | 31.3 ± 8.50^b^ | 36.0 ± 12.9 | 36.8 ± 12.3 | 24.5 ± 10.8^b^ | 0.7000 |
| 1L-13 | 23.6 ± 4.34 | 30.9 ± 4.47 | 32.3 ± 5.43 | 22.3 ± 5.17 | 0.3656 |
| 1L-15 | 127 ± 43.0 | 91.5 ± 17.4 | 129 ± 27.4 | 88.3 ± 25.8 | 0.8100 |
| IP-10 | 83.0 ± 9.08 | 95.9 ± 12.0 | 77.9 ± 6.94 | 92.2 ± 9.47 | 0.6890 |
| KC (CXCL1) | 894 ± 271 | 455 ± 132 | 332 ± 104 | 952 ± 337 | 0.2548 |
| LIF | 0.605 ± 0.390^b^ | 0.493 ± 0.091 | 0.790 ± 0.290 | 1.01 ± 0.297 | 0.4906 |
| M-CSF | 10.9 ± 4.08^b^ | 9.62 ± 2.68 | 6.52 ± 3.26^c^ | 12.7 ± 4.66 | 0.4937 |
| MIG | 214 ± 82.8 | 183 ± 34.8 | 160 ± 55.2 | 227 ± 70.0 | 0.7171 |
| MIP-1α | 36.1 ± 14.7^c^ | 69.9 ± 12.9^b^ | 41.7 ± 17.5^c^ | 56.5 ± 15.1 | 0.6285 |
| MIP-1β | 28.2 ± 13.4 | 62.8 ± 23.7 | 42.3 ± 11.8 | 36.0 ± 11.2 | 0.7930 |
| MIP-2 | 92.8 ± 19.9 | 69.5 ± 11.2 | 76.1 ± 10.2 | 75.9 ± 19.9^b^ | 0.6523 |
| RANTES | 36.2 ± 2.79 | 35.4 ± 4.48 | 30.4 ± 3.89 | 32.9 ± 6.18 | 0.6836 |
| TNFα | 20.0 ± 4.56 | 16.0 ± 3.23 | 11.3 ± 5.13 | 19.0 ± 4.96 | 0.5033 |
| VEGF | 0.360 ± 0.0525 | 0.483 ± 0.0541 | 0.423 ± 0.0428 | 0.466 ± 0.0584 | 0.4929 |

^a^By the Kruskal-Wallis test with Dunn's multiple comparisons test. A value of P <0.05 was considered

statistically significant.

^b^ One sample detected as out of range of the standard curve.

^c^ Two samples detected as out of range of the standard curve.

^d^ Three samples detected as out of range of the standard curve.

***Table S6. Related to Figure 6.*** Serum Hormones Levels of Muc2^-/-^ Mice Fed the MD (n=9), Olive Oil (n=10), Corn Oil (n=9) and Milk Fat Diets (n= 9).

| Serum Hormones  Mean ± SEM, by Diet | | | | | |
| --- | --- | --- | --- | --- | --- |
|  | MD  (n=9) | Olive Oil  (n=10) | Corn Oil  (n=9) | Milk Fat  (n=9) | *P* Value^a^ |
| Amylin | 54.8 ± 20.5 | 33.1 ± 6.64^b^ | 25.9 ± 5.38 | 30.3 ± 8.78 | 0.5328 |
| Ghrelin | 27.7 ± 14.5^c^ | 41.9 ± 19.0^c^ | 25.2 ± 10.8^c^ | 19.8 ± 11.2^c^ | 0.8745 |
| Peptide YY | 527 ± 51.0 | 628 ± 87.7^d^ | 499 ± 140 | 894 ± 265^d^ | 0.4159 |
| Glucagon | 81.8 ± 10.4 | 82.9 ± 13.8 | 73.4 ± 10.3 | 94.3 ± 10.3 | 0.6337 |
| C-Peptide | 2070 ± 286 | 1885 ± 168 | 2159 ± 196 | 2125 ± 269 | 0.9168 |
| Resistin | 29658 ± 3564 | 28744 ± 3940 | 31648 ± 2603 | 34710 ± 3793 | 0.6650 |
| Glucagon-Like Peptide 1 | 80.9 ± 22.5 | 96.4 ± 26.8^b^ | 49.9 ± 11.9^b^ | 90.8 ± 35.6^b^ | 0.7899 |

^a^ By the Kruskal-Wallis test with Dunn's multiple comparisons test. A value of P <0.05 was considered statistically significant.

^b^ One sample detected as out of range of the standard curve.

^c^ Four samples detected as out of range of the standard curve.

^d^ Two samples detected as out of range of the standard curve.

***Figure S6. Related to Figure 6.*** Type of fat did not influence total short chain fatty acids, butyric or propionic acid production in the cecum. By the Kruskal-Wallis test with Dunn's multiple comparisons test. A value of P <0.05 was considered statistically significant.

**A**

**B**

**C**

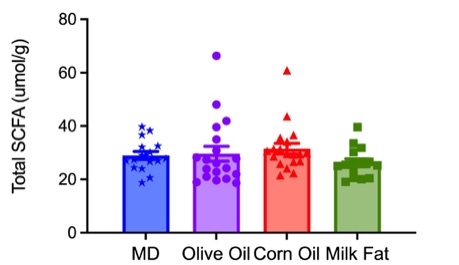

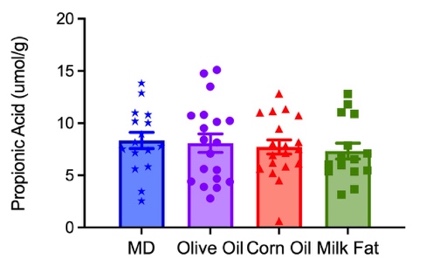

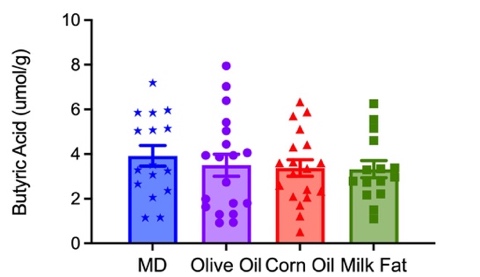

**Figure S7. Related to Figure 7.** Colon samples collected post-diet intervention and sequenced for 16S rRNA at the V4-V5 region. (A) Beta-diversity plots measured by robust CLR transform for cage batch and sex effects. (B) Alpha-diversity metrics for the colon show no statistic differences. (C) Beta-diversity statistics for Robust Aitchison (RPCA) pair-wise distances. (D) Songbird multinomial differential plot for ‘diet’. Data represent n=7-14 mice per group.

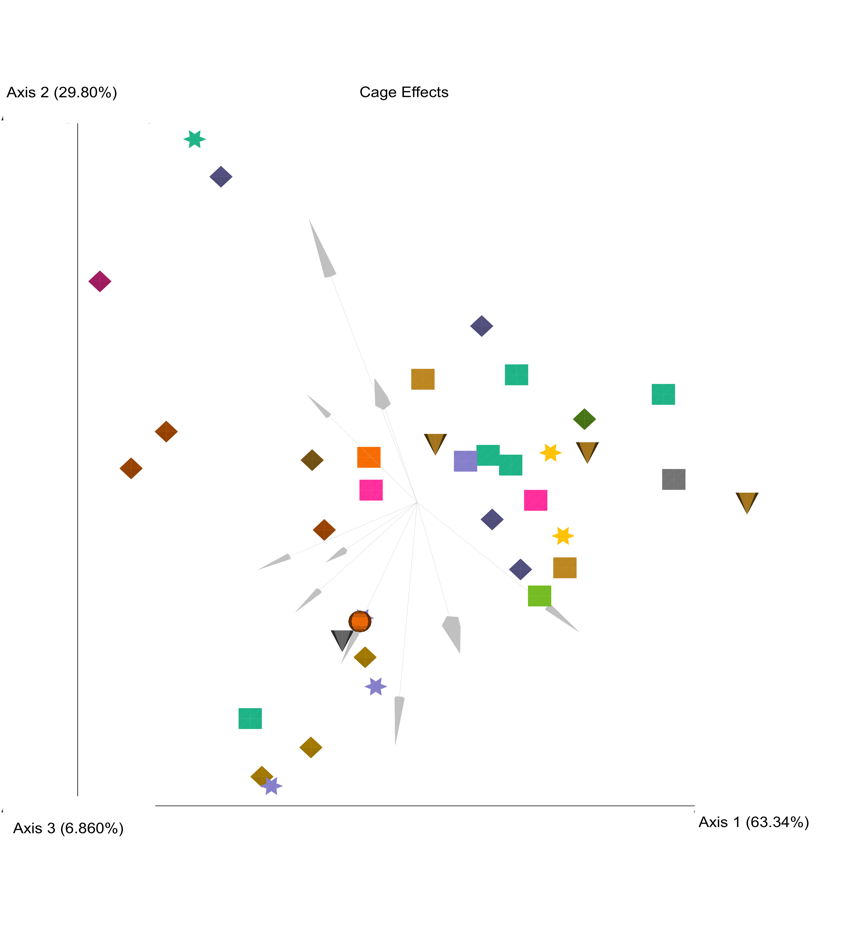

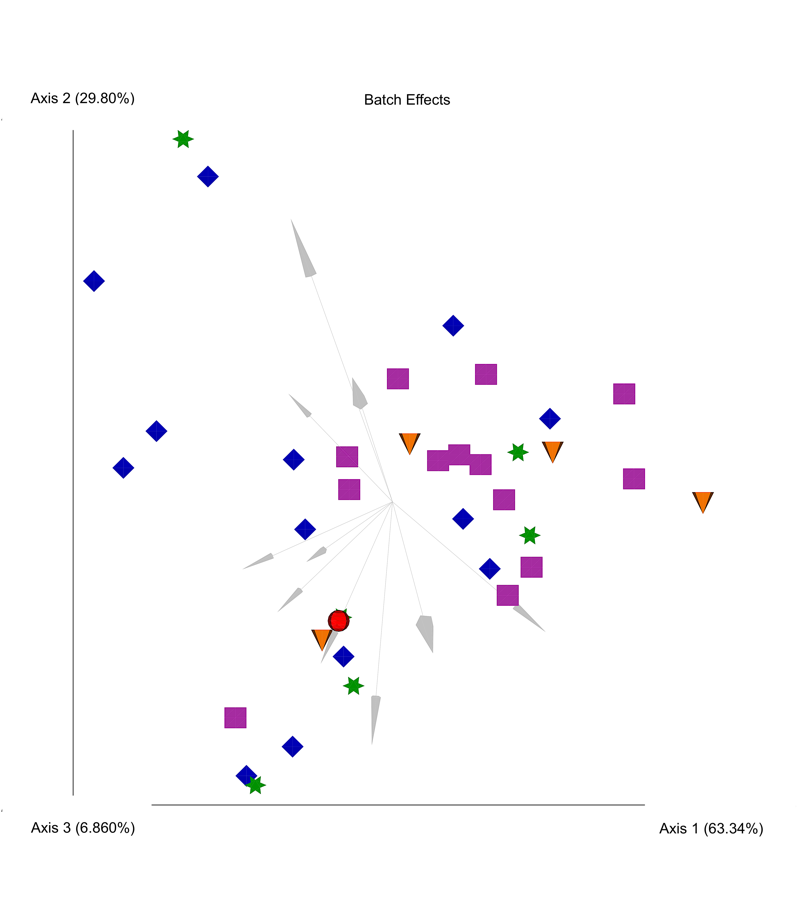

*Each shape and corresponding color represent a different cage or batch. Group effect was determined by the use of a PERMANOVA test on the robust CLR transformed data – no obvious clustering by cage or batch.

Cage Effects*

Batch Effects*

A

| Alpha-Diversity Metrics | P-value^a^ | H-value |
| --- | --- | --- |
| ASV Richness | 0.1604 | 5.1601 |
| Shannon Diversity Index | 0.3526 | 3.2648 |
| Faith’s Phylogenetic Diversity | 0.2955 | 3.7103 |
| Pielou’s Eveness | 0.3951 | 2.9774 |

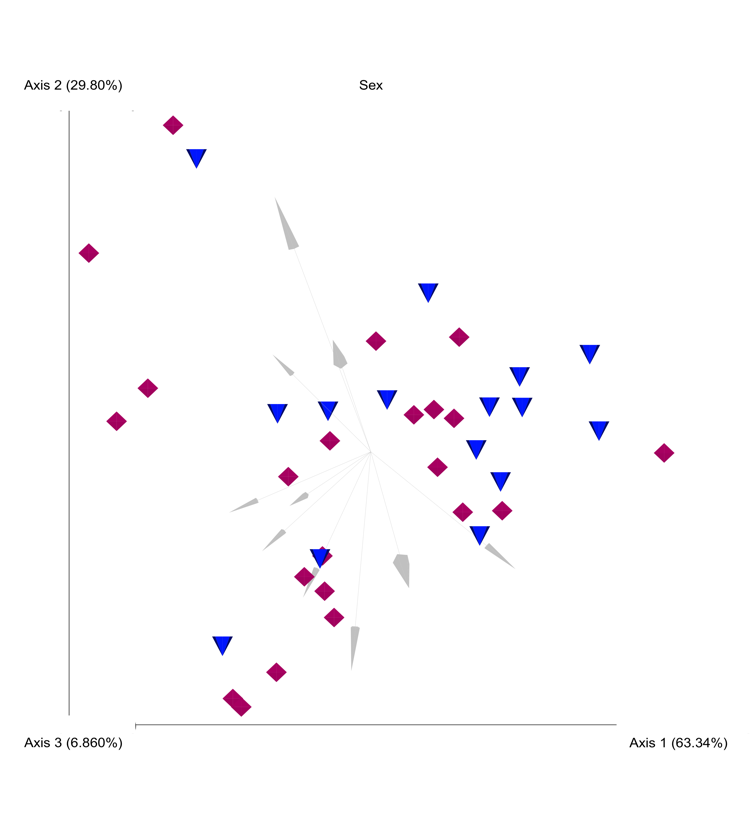

^a^Kruskal-Wallis test. A value of P <0.05 was considered statistically significant.

B

Sex*

*Diamonds = female, triangle = male. Analysis by robust

CLR transform – no obvious clustering by sex.

C

| Pairs | P-value | q-value | pseudo-F |
| --- | --- | --- | --- |
| CO - MD  CO - MF  CO - OO  MD - MF  MD - OO  MF - OO | 0.004  0.044  0.001  0.438  0.617  0.293 | 0.0120  0.0880  0.0060  0.5256  0.6170  0.4395 | 6.9317  3.5216  8.9532  0.8375  0.3664  1.1778 |

Pairwise PERMANOVA. A value of P <0.05 was considered statistically significant.

D

Songbird multinomial differential plot

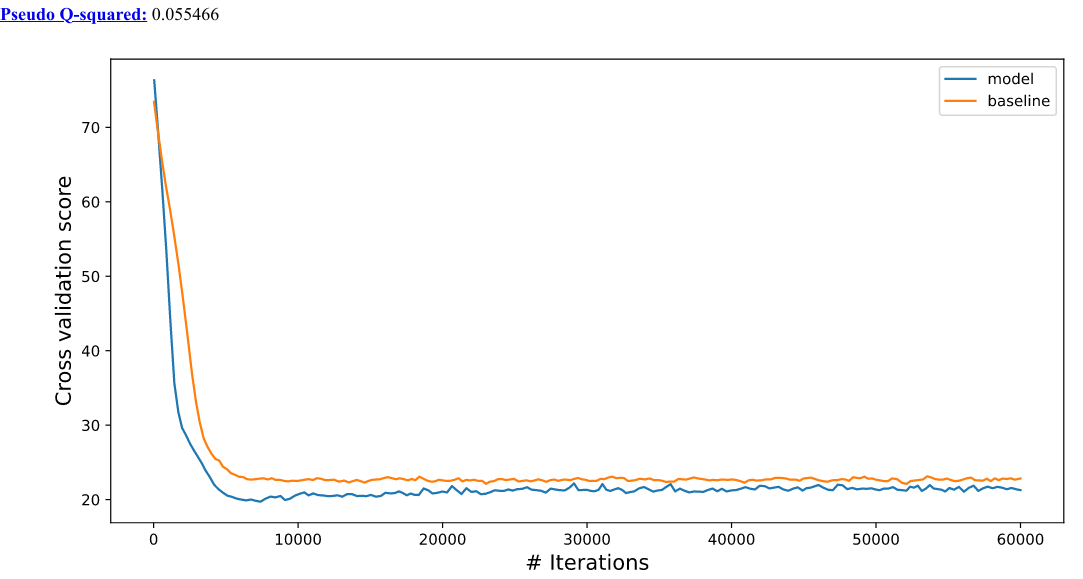
